# Multi-modal benchmarking of the Ultima UG100 and Illumina NovaSeq sequencing platforms using clinically relevant FFPE tissues

**DOI:** 10.64898/2026.03.11.710846

**Authors:** Quentin Bayard, Zelda Mariet, Anna Schaar, Ekaterina Rozhavskaya, Maria Mahfoud, Alex J. Cornish, Elo Madissoon, Fayssal Sadi-cherif, Rory Sinnott, Choiwai Maggie Chak, Almoatazbellah Youssef, Ramona Erber, Arndt Hartmann, Markus Eckstein, Silvia Lopez Lastra, Almudena Espin-Perez

## Abstract

Emerging high-throughput sequencing technologies promise lower costs and higher scalability, yet their performance on archival clinical samples remains poorly characterized. Here, we benchmarked Ultima Genomics UG100 against Illumina Novaseq platforms across single-nuclei RNA-seq (snRNA-seq), whole-transcriptome (WTS), whole-exome (WES), and whole-genome sequencing (WGS) using FFPE tissues from oncologic and immune-mediated diseases. Across matched samples, we systematically assessed data quality, coverage profiles, error spectra, variant concordance and transcriptomic reproducibility.

UG100 produced highly comparable results to Illumina, capturing key oncogenic and immune-related transcripts, accurately resolving cellular composition in snRNA-seq, and maintaining sensitivity for lowly expressed genes, despite characteristic insertion-biased indels and modest differences in multi-mapping reads. Discrepancies were subtle, largely limited to pseudogene and non-coding transcripts, and did not affect pathway-level conclusions. Ultima UG100 platform prioritized high precision and reduced low-frequency artifacts, offering a cleaner but more conservative variant-calling profile compared to the more sensitive, yet noisier, Illumina/DRAGEN workflow.

This multimodal, clinically oriented assessment provides the first comprehensive evaluation of UG100, demonstrating its translational utility in population-scale genomics, and highlighting the potential for emerging sequencing technologies to lower the cost of biomedical research and clinical diagnostics.

## Introduction

High-throughput sequencing technologies have revolutionized biomedical research, enabling high-resolution interrogation of genomes and transcriptomes across diverse disease contexts at the tissue and cellular level. Illumina’s sequencing-by-synthesis (SBS) technology has long set the industry standard, supporting population-scale studies - including whole-transcriptome (WTS), whole-exome (WES), whole-genome sequencing (WGS) - as well as single-cell or single-nucleus RNA sequencing (scRNA-seq/snRNA-seq). Illumina’s platforms have consistently delivered reliable, high-quality data and thus achieved broad adoption across research and diagnostic laboratories worldwide.

Today, the rapid expansion of AI-driven analyses in drug discovery, biomarker development, and clinical diagnostics has created unprecedented demand for large-scale, high-quality, multi-omic datasets. Accurate, reproducible, and scalable sequencing is now essential to train, validate, and deploy computational models that can extract actionable insights from complex biological systems. This growing demand has spurred innovation in alternative sequencing chemistries and platforms, including the recently published mostly natural sequencing-by-synthesis (mnSBS) and sequencing by expansion (SBX), which aim to reduce cost and increase throughput while maintaining sufficient accuracy for both variant- and transcript-level analysis ^1–5^.

Ultima Genomics’ sequencing platforms employ mnSBS, a novel sequencing chemistry which uses a low fraction of labeled nucleotides, combining the efficiency of non-terminating chemistry with the throughput and scalability of optical endpoint scanning within an open fluidics system. Initially implemented on the UG100 platform, this technology is reported to reduce whole-genome sequencing costs by over two-fold in a high-throughput setting versus the NovaSeq platform, and currently boasts a sequencing cost of less than $0.01 per cell for snRNA-seq. The recently introduced UG200 series builds upon this architecture through enhanced throughput, updated amplification workflows, and improved coverage uniformity, further extending the scalability of mnSBS. As such, mnSBS has the potential to scale biomedical data generation, a key component for advancing the field of precision medicine and drug discovery and development. Yet, comprehensive, cross-modality benchmarking against established SBS platforms remains limited, particularly on clinically relevant specimens such as formalin-fixed, paraffin-embedded (FFPE) tissues and in complex disease contexts, and is paramount to ensure the biological validity of mnSBS data and subsequent conclusions ^5–7^.

To address this, we conducted a comprehensive, multi-modal comparison of the Ultima UG100 and Illumina NovaSeq platforms using retrospectively collected FFPE specimens, spanning oncologic and immune-mediated disease pathologies, including diffuse large B-cell lymphoma (D), glioblastoma (G), muscle-invasive bladder cancer (M), ulcerative colitis (U), and Crohn’s disease (C). Samples were profiled across snRNA-seq, WTS, WES, and WGS, enabling a rigorous assessment of the UG100’s performance across a diverse range of analyses, from variant detection and transcript quantification to cellular composition and biological signal fidelity (Figure 1).

**Figure 1.**
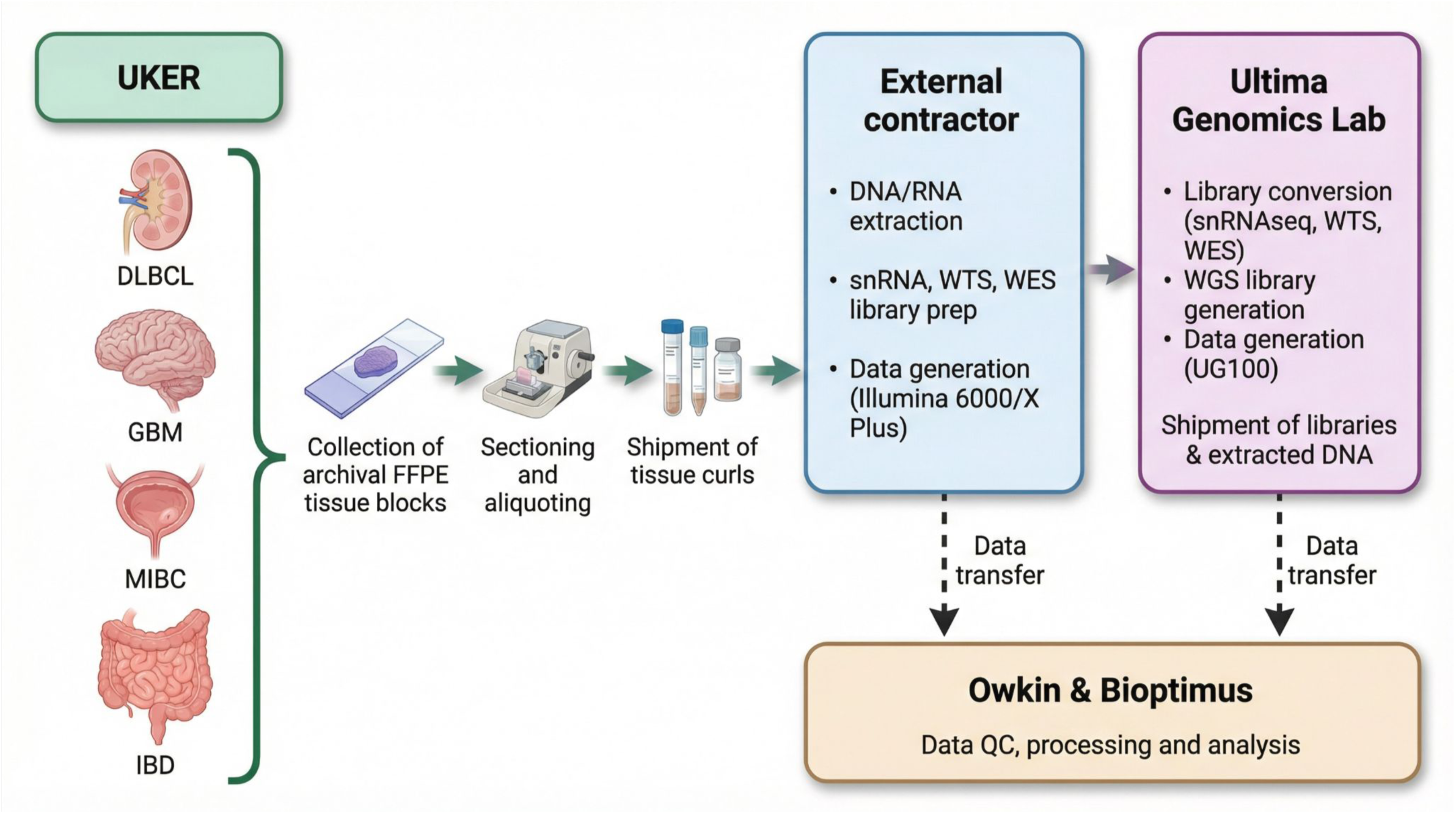
Sample preparation and multi-omics profiling. Three samples for each different indications were profiled using single-nuclei RNA sequencing (snRNAseq), whole transcriptome sequencing (WTS), whole exome sequencing (WES) and while genome sequencing (WGS) technologies in both UG100 and Illumina platforms. Diffuse large B-cell lymphoma (DLBCL), glioblastoma (GBM), muscle-invasive bladder cancer (MIBC), ulcerative colitis and and Crohn’s disease (IBD).

By systematically comparing sequencing metrics, coverage profiles, error spectra, variant concordance, and expression reproducibility under technology-optimized pipelines, we evaluate both the technical equivalence and practical utility of Ultima for translational and clinical research workflows. This study represents the first comprehensive, multi-modal evaluation of Ultima’s platforms on archival FFPE materials, and our findings provide the genomics community with an evidence-based perspective on its performance, limitations, and integration potential in the era of large-scale, AI-driven multi-omic analysis.

## RESULTS

### Sample preparation and multi-omics profiling

To enable a comprehensive and systematic benchmarking, we analyzed 15 FFPE samples spanning oncologic and inflammatory bowel disease (IBD) indications, aiming to evaluate technical performance, reproducibility, and biological concordance across two sequencing platforms while minimizing confounding from sample variability. Three patient derived tissues were profiled for each disease.

Samples were processed across multiple sequencing modalities: snRNA-seq and WTS for all specimens, WES for oncologic tissues (D, G and M), and WGS for IBD tissues (U and C) (Table 1). Samples were sequenced in parallel on the Illumina NovaSeq and Ultima UG100 platforms. One specimen per modality was sequenced in duplicate on each platform to assess intra-platform reproducibility (Methods).

DNA and RNA extraction, library preparation, and sequencing on the Illumina platform were performed by an external contractor (CeGaT, Germany), followed by library conversion and UG100 sequencing at Ultima Genomics Lab (Fremont, USA). WGS library construction and sequencing were conducted independently in the two facilities (Figure 1, Methods).

### Bulk RNA-seq benchmark

#### Sequencing metrics across platforms highlight differences across technologies and alignment pipelines

To benchmark the performance of UG100 relative to NovaSeq, we first looked into fundamental sequencing and alignment metrics. We observed distinct differences in insertion and deletion (indel) rates per base between the two sequencing platforms. As expected for SBS chemistry, Illumina reads exhibited a uniformly lower frequency of indel errors (Supplementary Figures 1A and 1B)^7^. The longer read lengths generated by Ultima (Supplementary Figure 1C) may also contribute modestly to these differences, as additional bases extend beyond the high-confidence alignment window. Importantly, no bias related to GC content was detected in either dataset (Supplementary Figure 1D).

In this FFPE cohort, UG100 produced reads averaging 120 base pairs (bp), shorter than the ∼ 250 bp described in early manuscripts (Supplementary Figure 1C), consistent with the higher degradation of FFPE samples processed here as compared to immortalized cell lines ^6^. Illumina NovaSeq reads averaged 90 bp, reflecting trimming of Illumina adapters from originally 101 bp(Methods).

UG100 libraries yielded substantially higher raw read counts compared to Illumina, which led to a proportional increase in duplicate reads consistent with sequencing saturation (Supplementary Figures 1E and 1F). To ensure a fair, head-to-head comparison between platforms, we downsampled the UG100 dataset to match the total read counts of the corresponding Illumina libraries (Methods). This normalization allowed us to directly compare metrics such as coverage uniformity, variant detection and gene quantification without confounding effects from differing sequencing depths.

On all reads, the Ultima dataset contains slightly fewer unmapped and multi-mapped reads than Illumina and slightly more uniquely mapped reads (Figure 2A). The Ultima dataset also contains a slightly higher proportion of high-quality reads than Illumina (Supplementary Figure 2A), although less high-quality reads in total numbers (Supplementary Figure 2B). However, we identified none of these differences to be significant. We hypothesized that Illumina reads are mapping to similar sequences elsewhere in the genome (pseudogenes or members of the same family) due to shorter read length. For the reads that are well mapped, Ultima contains a higher proportion of spliced reads than Illumina, but not significantly so (Supplementary Figure 2C).

**Figure 2.**
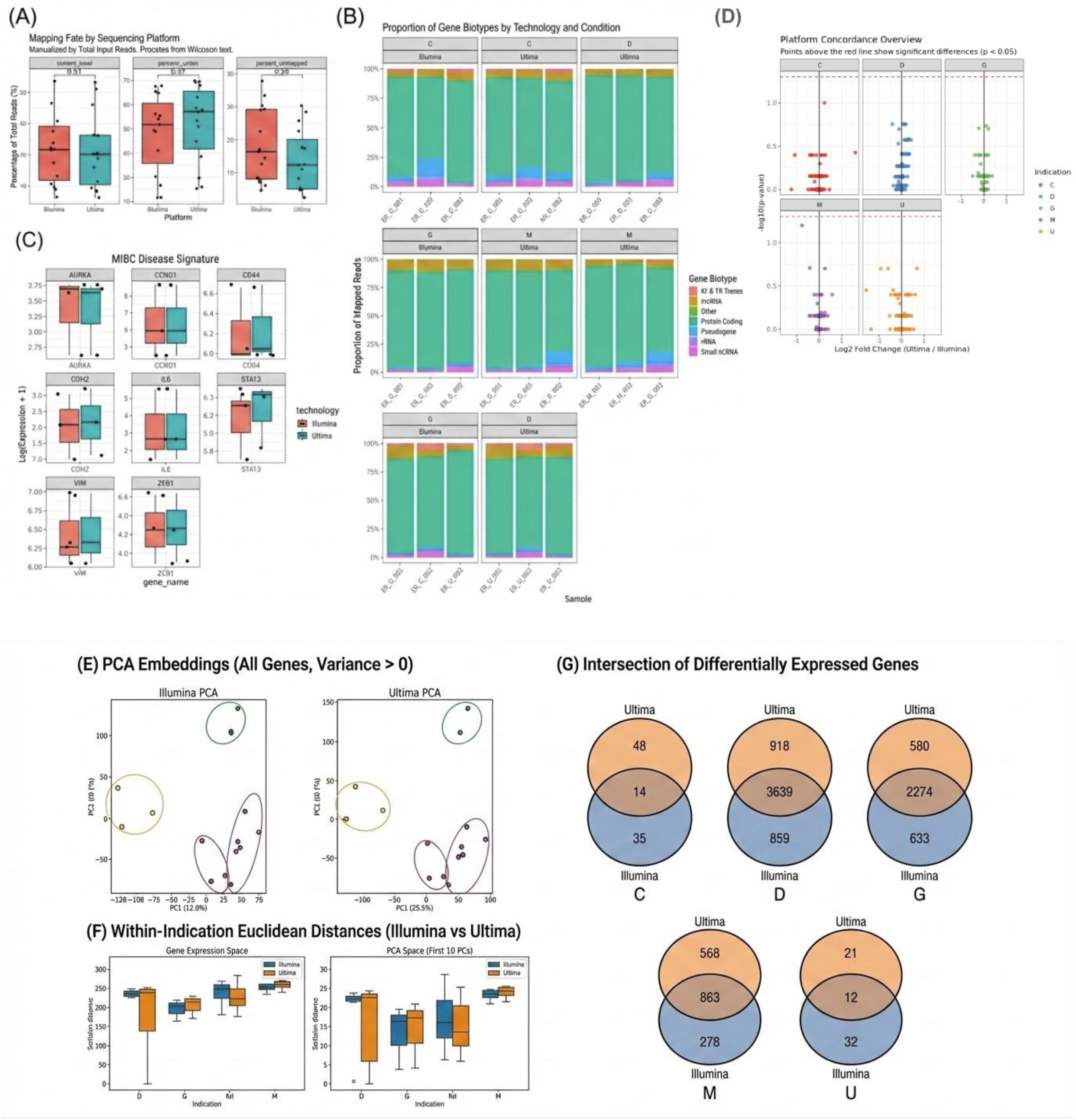
Sequencing metrics, biotypes and biology assessment. **(A)** Proportion of uniquely mapped, multi mapped and unmapped reads **(C)** Biotype representation across indications and technologies **(C)** Genes expression from selected MIBC signature and **(D)** Log fold change between Ultima and Illumina gene expression of all selected markers. **(E)** Clustering of samples from PCA embeddings using VST normalized data across all genes with variance > 0 in Illumina and UG100 technologies **(F)** Euclidean distance between samples from the same disease in the common embeddings (VST normalization with batch correction) **(G)** Intersection of differentially expressed genes between each disease and the rest of diseases obtained from Illumina and Ultima UG100 .

Finally, to assess reproducibility, we compared counts per gene from technical replicates of consecutive sections. Both Illumina and UG100 demonstrated high correlation (r = 0.95 and 0.96, respectively; Supplementary Figure 2D) indicating excellent reproducibility even from FFPE samples. Minor variability likely reflects biases in library preparation or sequence composition.

Overall, these results confirm that, despite platform-specific differences in indel profiles and read characteristics, both technologies provide consistent quantification suitable for reliable expression analyses.

#### Ultima and Illumina show similar gene expression levels of selected disease and biomarker genes

We observed that Illumina detected a slightly higher percentage of genes than Ultima (Supplementary Figure 3A) (p-value < 0.001), prompting a closer examination of gene biotypes and read alignment characteristics. Gene biotype, as defined by Ensembl, classifies transcripts into categories such as protein coding, pseudogene or long non-coding RNA ^8^. We identified more pseudogenes (∼0.23%) and lncRNA transcripts (∼0.11%) and fewer miRNA (∼0.12%) in non expressed genes from Ultima than from Illumina, although the findings were non-significant (Supplementary Figure 3B). These differences are consistent with platform-specific technical features: shorter reads in Illumina can be flagged as multimappers since they can map to both protein coding and closely related pseudogenes, while Illumina’s highly optimized adapter chemistry may enhance capture of short RNAs. While these differences were not substantial, they highlight platform-specific biases inherent to sequencing chemistry and read mapping near the limit of detection.

To contextualize the biological relevance of these discrepancies, we examined closely the gene set detected by Illumina but absent in Ultima across all patients. Several of them showed high expression, including *LUZP4* (a cancer-testis antigen), *TDGF1* (Cripto-1, a well-studied oncogene), *TNP1* (a candidate biomarker for testicular germ-cell tumours), *OR51Q1* (reported in pancreatic squamous cell carcinoma) and *LIMS3* (associated with immune response signatures)^9–13^ (Table 2). We also observed relevant oncologic and immunological genes in Ultima which were absent in Illumina, to a lesser extent (921 genes detected in Ultima and 2150 in Illumina). The distribution of these genes among biotypes confirms Illumina’s higher capacity to capture pseudogenes (Supplementary Figure 3C). Although these genes represented a small subset, their involvement in immunology and oncology pathways underscores the importance of platform-specific validation to ensure reliable biomarker identification and clinical interpretation. Interestingly, despite the discrepancies described above, both platforms captured similar percentages (∼15%) of low expressed genes (< 10 counts), indicating comparable sensitivity for rare transcripts (Supplementary Figure 3D).

Overall, biotype representation was similar between platforms but varied across disease indications and patients, as expected (Figure 2B). Oncologic samples were enriched for protein-coding genes, while immunoglobulins and T cell receptor genes were more abundant in IBD samples, independent of the sequencing technology ^14,15^ .

We hypothesize that, while Illumina’s higher base-call accuracy and error profile result in greater per-base accuracy, the extended read lengths by the Ultima platform provide greater genome spatial context. This increased context may overcome indel-related noise, leading to higher unique mapping rates and more definitive assignment to high-homology regions like pseudogenes. In contrast, the UG100 pipeline may attribute these same ambiguous reads to protein-coding genes, where the alignment confidence is relatively higher according to its scoring model. Consequently, UG100 datasets displayed an increased expression of protein-coding genes at the expense of other biotypes, whereas in Illumina reads were distributed more evenly across highly similar loci. These findings underscore the impact of platform-specific alignment behavior on observed gene biotype distributions and highlight the importance of considering read assignment confidence when interpreting comparative transcriptomic data.

To further assess platform performance, we evaluated structurally challenging genes, including those with high GC content, multiple isoforms, or repetitive regions (e.g., transcription factors, histones, and housekeeping genes) (Table 3). Across these gene categories, expression levels were highly comparable between platforms, suggesting comparable performance in technically demanding regions (Figure 2D). We then assessed the biological and clinical relevance of the observed expression patterns by examining key oncogenic drivers (e.g. TP53, MYC), markers of proliferation and differentiation (e.g., KRT5, GATA3, CCND1), immune signaling mediators (e.g., IL6, STAT3, CXCL8), and disease-specific biomarkers across oncologic and immune-mediated disease contexts. Expression was highly consistent between the platforms, demonstrating that both technologies reliably capture meaningful known transcriptional signals for downstream analyses and interpretation (Figures 2D, Supplementary Figures 4 A-H).

#### Ultima and Illumina recover expected biology and data can be combined without batch correction techniques

Samples from the same disease clustered closely within each sequencing platform, both before and after batch correction, indicating that both technologies robustly capture underlying biological variation (Figure 2E, Supplementary Figure 5A). Without correction, Ultima displayed slightly tighter clustering of samples within the same indication compared to Illumina, suggesting higher intrinsic consistency in gene expression profiles. Following batch correction, with sequencing technology specified as the batch variable and disease indication preserved as a biological covariate, Illumina samples showed a marked improvement in intra-indication clustering, surpassing Ultima in oncology datasets (Figure 2F). In contrast, Ultima maintained stronger intra-group coherence in IBD samples, even prior to correction (Supplementary Figure 5B). This differential effect suggests that while Illumina data contain residual batch-associated variance that can be effectively mitigated computationally, Ultima data exhibit greater inherent stability across sequencing runs and conditions. We hypothesize that immune-related transcripts in IBD (often expressed at a wide dynamic range) are more sensitive to amplification and adapter-related bias. Therefore, Ultima’s chemistry may capture them more accurately, while Illumina’s superior mapping of complex regions (often present in oncology) may allow the biological signals of the tumor to emerge upon removal of linear and correctable batch effects. Thus, the observed differences are likely due to a combination of sequencing technology biases and the underlying biology of the two disease types.

Differential expression analyses revealed differences in the number of genes identified by each platform when comparing individual disease groups to the remaining samples (Figure 2G). However, these discrepancies did not translate into meaningful differences at the pathway level. Only four pathways were uniquely detected by one technology, compared to 2,750 pathways shared between platforms (Table 4), and biotype composition of uniquely identified genes was comparable. Moreover, differences in differentially expressed gene detection were not attributable to gene length (Supplementary Figure 6A). These findings indicate that despite modest platform-specific differences in individual gene detection, both technologies converge on the same higher-order biological insights.

To assess systematic expression biases independent of disease context, we compared aggregated expression profiles across all samples. We identified 2,021 genes downregulated and 1,049 upregulated in UG100 relative to Illumina, corresponding to distinct but biologically coherent pathway enrichments (Table 5, Supplementary Figure 6B). Upregulated genes were enriched for protein-coding biotypes, whereas downregulated genes included a higher proportion of pseudogene, snRNA, and misc_RNA biotypes (Supplementary Figure 6C). This pattern may reflect the differences in alignment confidence and read assignment strategies between platforms: Illumina’s base quality may resolve ambiguous reads across homologous regions more confidently, assigning them to the pseudogene loci, while reads from Ultima may assign to the corresponding protein-coding gene instead, leading to apparent shifts in biotype representation.

Overall, biotype distributions were broadly consistent across the technologies but varied as expected by disease indication. Oncological samples displayed a higher representation of protein-coding genes, whereas IBD samples exhibited a greater abundance of immunoglobulin and T cell receptor transcripts, independent of sequencing technology (Figure 2B). Together, these results demonstrate that both UG100 and Illumina platforms capture biologically meaningful transcriptional variation with high fidelity, while differing in the nature of sequences that leads to subtle differences in gene biotype assignment and retaining biological signal after batch correction.

### Single-nuclei RNA-seq benchmark

#### Ultima and Illumina show similar relative change in cell counts when filtering out low quality cells

To benchmark the performance of the UG100 sequencing platform for fixed snRNA-seq data, we generated libraries from FFPE oncology and IBD samples using Single-Cell RNA Sequencing Flex (SCR Flex) for Illumina sequencing, and part of these were subsequently converted into Ultima libraries for sequencing (Methods). In our study, Ultima UG100 libraries initially had substantially higher sequencing depth than Illumina libraries; however, this increased depth did not appear to affect the overall data quality. Despite an average 5-fold increase in reads per cell between Ultima and Illumina, the median number of UMIs per cell was not substantially increased (Figure 3A), indicating that the additional reads largely represent oversampling rather than capturing the additional transcript diversity. Only in DLBCL did all three patients have higher UMI counts with Ultima. We performed quality control on each sample independently using automatic thresholding based on the distribution of gene counts and percentage of mitochondrial counts per cell (Figure 3B) as well as doublet detection (Figure 3C). This analysis and QC filtering approach resulted in comparable total fractions of low-quality cells detected for both technologies (Figure 3D). After filtering, Ultima datasets still maintained a higher sequencing depth, indicating the increased depth represents an overall shift in the gene expression distribution, but has no immediate effect on data quality itself (Methods).

**Figure 3.**
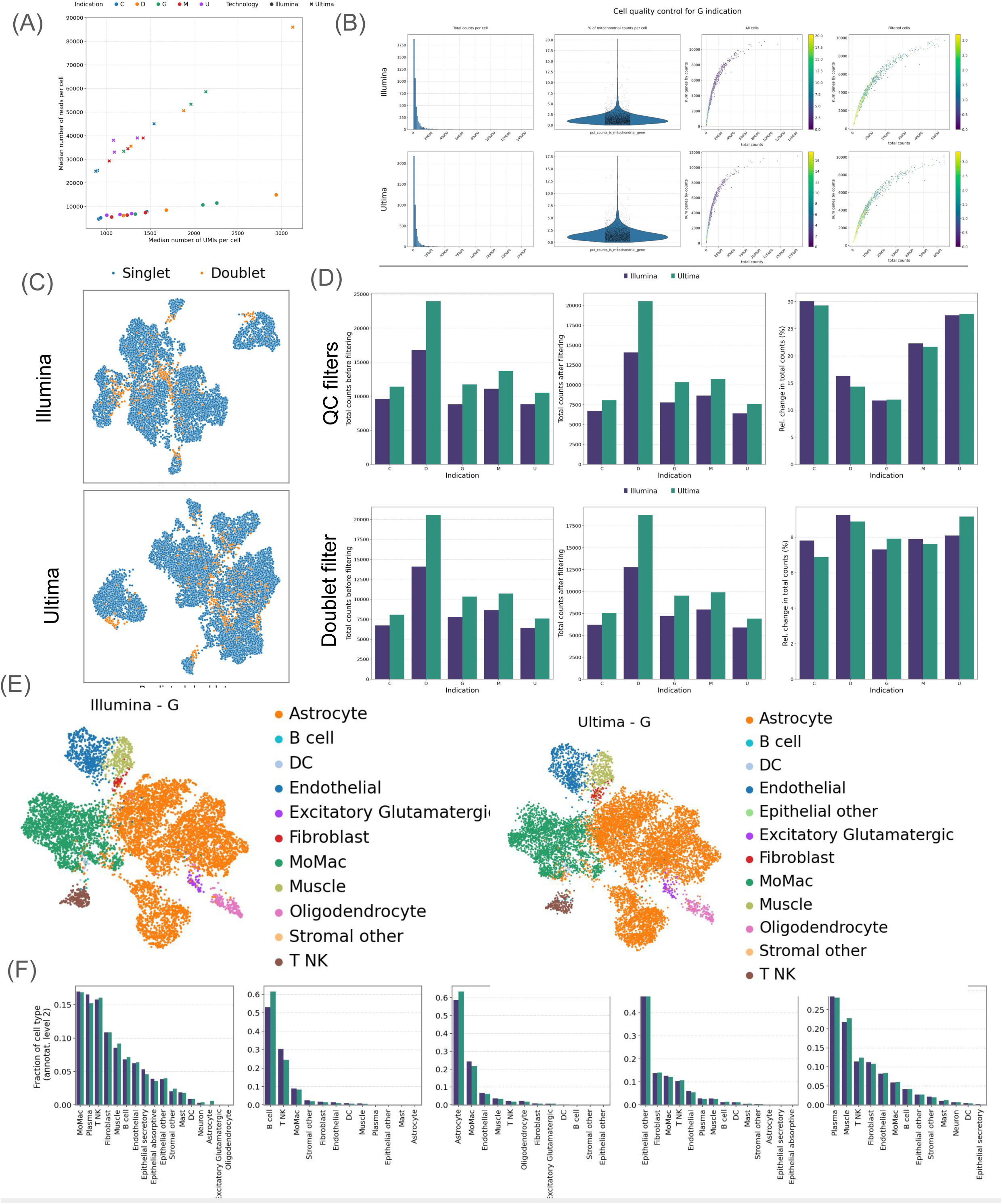
Single-cell RNA-seq comparison. **(A)** Number of UMIs and number of reads per cell (**B)** Comparison of QC for Illumina and Ultima UG100 on one sample for indication G **(C)** Doublet detection for Illumina (7.31%) and Ultima (7.91%) on indication G **(D)** Relative change in library size after filtering by gene and mitochondrial counts (top) and doublet detection (bottom) **(E)** UMAP embedding of all G sample in Illumina (left) versus UG100 (right) with cell type level 2 superimposed. **(F)** Fractions of cell types at level 2 for both technologies

#### Clustering and cell-type annotations are qualitatively similar

To further assess the impact of gene annotation and expression level differences on the cell-type interpretation, we performed hierarchical cell annotations (at multiple levels of granularity) across both platforms and disease indications using scANVI (Methods). When embedding cells from both technologies into a lower-dimensional UMAP representation, cells clustered primarily by biological cell type rather than by the sequencing technology, indicating a minimal or low technical variation (Figure 3E). This finding suggests that additional batch-effect correction was not required and that relevant biological variation was well-preserved. Both platforms recovered the major expected cell types for each disease indication and cells remained comparably well-integrated within each disease cohort.

Furthermore, the two platforms showed a high concordance in the proportions of each cell type across samples and annotation levels for the MIBC, GBM and IBD samples (Figure 3F, Methods). For DLBCL, we observed minor differences in cell capture within the B cell compartment. Statistically significant platform-specific deviations were observed only in the DLBCL cohort, and these deviations emerged progressively across annotation levels 2, 3, and 4. At level 2, Ultima detected a higher proportion of B cells that were most abundant, whereas Illumina captured more of the other abundant cell types out of which the second most abundant, T/NK cells, was significant. At level 3, this pattern refined into increased B-cell abundance in Ultima, contrasted with the elevated CD4⁺ T-cell proportions in Illumina. At the most granular level 4, the primary divergence centered on the malignant compartment: malignant B cells were more abundant in Ultima, while Illumina retained a comparatively larger fraction of T-cell populations. The enrichment of malignant B-cells could be explained by the higher number of captured UMI-s by Ultima in DLBCL.

### Whole Exome Sequencing benchmark

We evaluated the performance of the two platforms for WES using oncology samples and assessed multiple quality, mapping, and variant calling metrics. Unless otherwise stated, comparisons are reported at the level of platform–pipeline combinations rather than the sequencing platforms alone. Illumina data were processed using the DRAGEN pipeline, whereas Ultima data were processed using DeepVariant, reflecting the use of platform-optimized, best-practice analysis workflows for each technology. Accordingly, the reported differences primarily capture combined effects of sequencing and downstream processing rather than the sequencing chemistry alone.

#### Comparable sequencing quality and coverage, with minor, predictable platform-specific indel differences in low-complexity regions

Both technologies achieved high-quality data across all metrics. Mean target coverage was 130x for Ultima UG100 and 120x for Illumina, with over 95% of targeted bases covered at ≥20× in both platforms. Coverage uniformity across exonic targets was comparable, although Illumina exhibited slightly more consistent representation in the regions of extreme GC content. On-target rates were high for both datasets (93-95% for Ultima and 94-96% for Illumina) indicating an efficient capture and a minimal off-target sequencing.

Duplication rates were modestly higher for Ultima (mean 40-50% for Ultima vs. 35-45% for Illumina), resulting in a minor reduction in effective unique coverage, but the overall sequencing depth compensated for this effect. Base quality distributions showed that over 90% of Ultima reads and 94% of Illumina reads exceeded Q30, reflecting excellent per-base accuracy for both platforms. Notably, Illumina reads demonstrated slightly lower Q-scores at the 3′ read ends, consistent with the platform’s paired-end design.

Illumina exhibited a uniformly low frequency of indel errors across genomic contexts, as observed in the other modalities. Exploration of local sequence complexity using Shannon entropy showed that Ultima indels occurred in regions with significantly reduced sequence complexity compared to Illumina: the mean entropy of 50-bp flanking regions was 1.792 for Ultima versus 1.899 for Illumina indels. This difference was highly statistically significant (Mann–Whitney U test, one-sided, p < 0.001) and corresponds to a Δ entropy of 0.107 (∼5.6% lower entropy for Ultima). The reduced entropy reflects increased repetitiveness in the surrounding sequence, including homopolymers and short tandem repeats, which indicates that Ultima indel errors are systematically biased toward lower-complexity genomic regions (Figures 4A and 4B).

**Figure 4.**
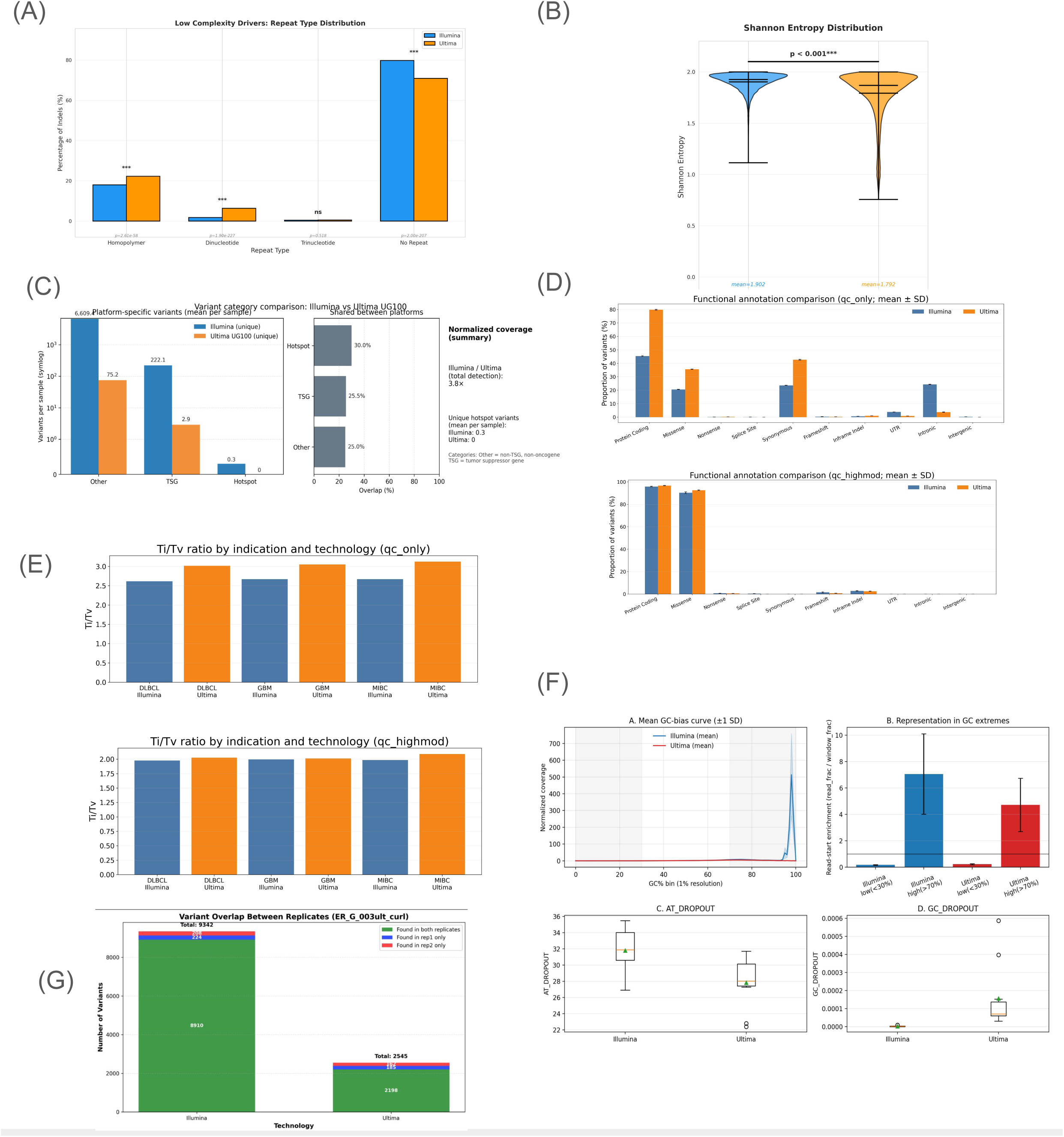
WES comparison. **(A)** Distribution of indel errors in WES data **(B)** Shannon entropy distribution of indels in WES data **(C)** Illumina vs Ultima platform-specific (unique) variants per sample, overlap (%) shared between platforms for the same categories (25.0%, 25.5%, 30.0%) and call out with the 3.8× Illumina/Ultima total detection ratio and hotspot uniqueness callout. **(D)** Functional annotation comparison for QC only (variants that pass VCF call-quality filtering, no functional/impact restriction. It includes IMPACT = HIGH, MODERATE, LOW, MODIFIER as present in CSQ) and QC high mode (qc_only filtering and additionally filter for only variants with VEP IMPACT ∈ {HIGH, MODERATE} (from the CSQ annotation) **(E)** Transition/transversion (Ti/Tv) ratio by indication and technology for qc and qchigh mode as defined in D. **(F)** GC content analysis for mean values, representation in GC extremes, in low-GC / AT-rich regions (AT_DROPOUT) and in high-GC regions GC_DROPOUT (**G**) Variant overlap in the GBM replicated sample

Overall, these metrics demonstrate that Ultima UG100 achieves sequencing and mapping quality equivalent to Illumina WES for most parameters, with small and predictable differences reflecting the distinct sequencing chemistries and read structures of each platform. The additional enrichment of Ultima indels in low-complexity regions highlights a characteristic, context-dependent error mode that should be considered in downstream variant calling. Nonetheless, the overall performance of both platforms provides a strong foundation for robust comparative genomic analyses.

#### Robust detection of oncogenic hotspot across platforms and pipelines

To ensure that differences in variant calling sensitivity were not driven by the higher raw sequencing depth of the Ultima platform (130x versus. 120x mean coverage), Ultima datasets were downsampled to match Illumina mean target coverage prior to variant discovery (Methods).

When examining variant categories at normalized coverage, Illumina detected an average of 6,609.4 “other” (non-tumor suppressor gene [TSG], non-oncogene) variants, 222.1 TSG variants, and 0.3 known hotspot variants per sample that were unique to this technology. In contrast, Ultima UG100 detected far fewer platform-specific variants, with averages of 75.2 “other” variants, 2.9 TSG variants, and no hotspot variants per sample (Figure 4C). This resulted in an Illumina-to-Ultima ratio of 3.8x for total variant detection. The overall overlap between technologies remained modest across categories, with 25.0% of “other” variants, 25.5% of TSG variants, and 30.0% of hotspot variants shared between platforms.

BAM inspection in IGV revealed that nearly all variants were present in reads from both platforms, indicating that discrepancies primarily arise from differences in bioinformatic pipelines. Specifically, Illumina data processed through DRAGEN generated a larger number of low-confidence calls, whereas Ultima UG100 read analyzed with DeepVariant, optimized for this platform, applied more stringent calling thresholds. Variants classified as “somatic” in Illumina but not called in Ultima were frequently labeled “RefCall” or “LowCall” by the latter caller (Supplementary Figure S7).

Importantly, recurrent and clinically actionable driver mutations were consistently detected by both platforms. For example, activating mutations in EZH2 and PTEN loss-of-function alterations are known to be recurrent in DBLCL and GBM patients, respectively ^16–18^ . Similarly in mesothelioma patients, CDKN2A, PIK3CA and TP53 mutations are known to be drivers of this cancer development ^17,19^. These observations confirm that clinically actionable variants are reliably captured (Table 6).

Aggregated analysis showed an average of 78.1 Ultima-only calls per sample across all categories, compared with 6,831.8 Illumina-only calls. This ∼87-fold difference in unique calls suggests that the Illumina/DRAGEN pipeline adopts a significantly more permissive calling posture compared to the Ultima/DeepVariant approach. While a lower number of unique calls can be consistent with higher specificity, these differences likely reflect the use of distinct alignment and variant-calling pipelines rather than sequencing platform performance alone.

Variant allele frequency (VAF) analysis supported this interpretation: Illumina-unique variants were enriched at low-VAF (<10%), consistent with low-confidence or borderline calls, whereas Ultima-only variants had a higher VAF, reflecting a more conservative calling approach. Manual inspection of sequences corresponding to some Illumina-unique variants showed identical underlying sequences in both datasets, further indicating that the differences are driven by pipeline stringency rather than base divergence. Importantly, the 30.0% overlap in hotspot variants and the 25.5% overlap in TSG variants demonstrate that despite the massive difference in “background” calls, both platforms converge on the most biologically significant regions of the exome.

Across all oncology indications (DLBCL, GBM, and MIBC), we compared Illumina and Ultima WES variant callsets derived from the same pipeline-produced VEP-annotated VCFs. On the QC-only callset, Illumina yielded more variants per sample on average (40167 vs 21069). Cross-technology concordance was modest (mean Jaccard index = 0.50) with shared variants representing 51% of Illumina calls and 96% of Ultima calls. This asymmetry indicates that the majority of Ultima calls were also detected by Illumina, whereas Illumina included a substantial number of additional platform-specific variants.

Transition/transversion (Ti/Tv) ratios computed on QC-only single-nucleotide variants (SNVs) were 2.65 for Illumina and 3.06 for Ultima. Restricting to higher-impact calls, reduced callset sizes and shifted functional composition toward protein-coding consequences, with mean protein-coding proportions of 95.8% for Illumina and 96.6% for Ultima (Figure 4D, Table 7). Under this high/moderate-impact subset, Ti/Tv ratios converged (1.99 Illumina; 2.04 Ultima) (Figure 4E).

Using Picard GC-bias profiling, we also quantified the GC-dependent coverage distortion and dropout for exome targets generated on Ultima and Illumina. Across 1%-wide GC bins spanning 0–100% GC, both technologies exhibited broadly similar GC-biased shapes characterized by the depletion in low-GC bins and the enrichment in high-GC bins. When collapsing bins into three strata (<30%, 30–70%, >70% GC), normalized coverage in the low-GC windows was reduced for both platforms and was higher on Ultima than Illumina (mean normalized coverage 0.231 vs 0.173), whereas high-GC windows showed a strong enrichment on both platforms and was higher on Illumina than Ultima (7.06 vs 4.72). Consistent with these distortions, the fraction of read starts mapping to GC extremes differed by platform: Illumina showed a higher representation in >70% GC (3.90% vs 2.61%), while Ultima showed a higher representation in <30% GC (3.04% vs 2.28%). Dropout metrics further supported platform-specific differences: AT dropout was lower on Ultima than Illumina (mean 27.82 vs 31.81; Mann–Whitney U p=0.021), whereas GC dropout was higher on Ultima than on Illumina (mean 1.6×10-4 vs 3.0×10−6 ; Mann–Whitney p=1.7×10−4), although GC-dropout values were small in absolute terms for both platforms (Figure 4F).

Finally, cross-sample reproducibility was evaluated by calculating the per-variant concordance across replicates. Shared variants demonstrated 95.4% reproducibility in Illumina and 86.4% in Ultima, further reinforcing the consistent variant detection within each platform’s optimized pipeline. The higher replicate concordance observed for Illumina is consistent with its greater sensitivity and inclusion of low-confidence calls (Figure 4G).

Collectively, these analyses indicate that the primary source of discordance between platforms arises from differences in variant-calling stringency and bioinformatic configuration rather than from underlying sequencing chemistry. Ultima UG100 produces a more conservative callset with high overlap for biologically and clinically relevant variants, while Illumina/DRAGEN generates a broader variant landscape that includes a larger fraction of low-VAF and low-confidence calls. Importantly, both platforms reliably detect key oncogenic and tumor suppressor alterations, supporting their suitability for translational exome profiling when interpreted within their respective bioinformatic frameworks.

### Whole Genome Sequencing benchmark

We next assessed the performance of both platforms for generating WGS data by analyzing germline IBD samples (Methods), focusing on variant detection, concordance and platform-specific calling characteristics. As global sequencing quality and coverage metrics closely mirrored those described earlier in our WES analysis, we concentrated our WGS assessment on downstream variant level comparisons. Throughout this section, references to “Illumina” and “Ultima” denote the Illumina+DRAGEN and Ultima+DeepVariant workflows, respectively.

#### Germline indel detection shows comparable sensitivity with differential false-positive burden

To benchmark indel performance, we compared each platform’s indel calls against the Genome in a Bottle (GIAB) truth set for sample HG001 (NA12878) on GRCh38, which contains 571,515 high-confidence indels ^20^. Ultima recovered 490,019 true indels (85.74% sensitivity) with 20,972,845 false positives (2.28% precision), while Illumina recovered 482,111 true indels (84.36% sensitivity) with 6,607,238 false positives (6.80% precision).

Overall, UG100 showed slightly higher sensitivity but a substantially larger false-positive burden (3.17x more FPs), whereas Illumina achieved higher precision (2.98x higher), yielding a higher F1 score (0.1259 vs 0.0445) (Figure 5A left). These results indicate that the Ultima/DeepVariant workflow, under the applied configuration, favors recall over precision in indel calling relative to Illumina/DRAGEN. However, UG100 sensitivity dropped markedly after downsampling (to ∼0.097), while precision increased (to ∼0.315). Importantly, this change should be interpreted in the context of both reduced sequencing depth and a non-identical callset stage/representation of the downsampled VCFs, which can shift the precision–recall balance independently of sequencing chemistry (Figure 5A right).

**Figure 5.**
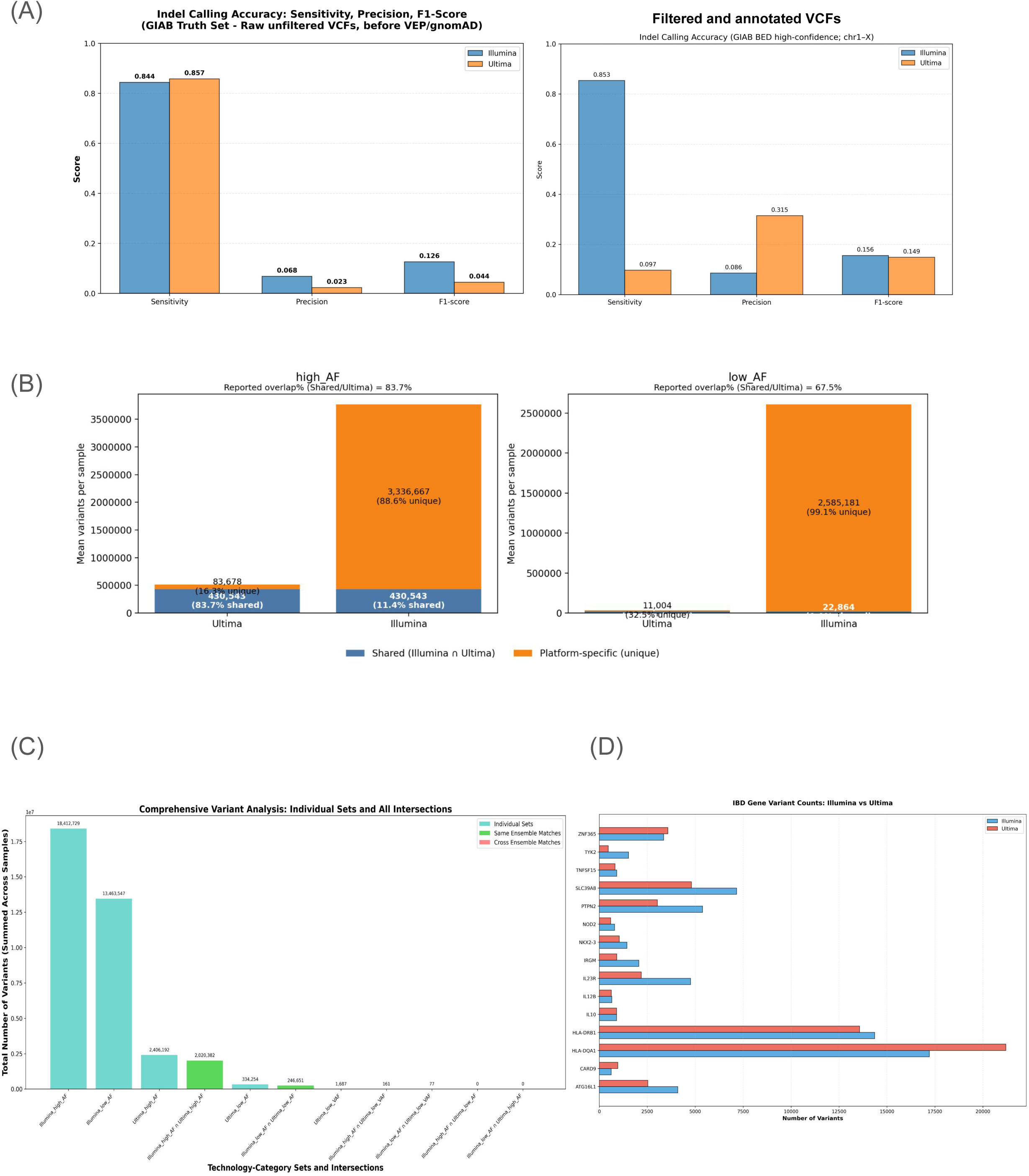
WGS comparison. **(A)** Shared high and low AF variants between technologies **(B)** Total number of variants per AF and VAF category **(C)** Number of IBD variants per selected gene of interest **(D)** Indell calling accuracy between both platforms in the GIAB data set.

#### Comparable germline variant detection with workflow-dependent sensitivity and low-frequency call differences

Notably, while the GIAB benchmarking for HG001 utilized high-depth datasets for both platforms to assess peak performance, all subsequent comparative analyses of the IBD cohort were performed using standardized, equal-coverage downsampled reads to ensure a fair real-world comparison. We examined germline variant detection, stratifying calls by population allele frequency (AF) and variant allele frequency (VAF) to distinguish high-confidence germinal variants from potential artifacts. For high AF variants, representing true germline events, Ultima UG100 detected an average of 514,221 variants per sample, compared to 3,767,210 variants per sample with Illumina, corresponding to a Ultima/Illumina ratio of 0.14x. In contrast, low AF variants, representing likely artifacts, were detected at higher rates by Illumina (2,608,045 variants per sample) than Ultima (33,868 variants per sample). Even when accounting for Illumina’s higher total call volume, low-frequency artifacts constituted ∼70% of the Illumina call set compared to only ∼6% of Ultima’s, representing a disproportionate enrichment of noise in the Illumina workflow at these specific filter settings.

Overlap analysis demonstrated high concordance for high AF variants (83.7%, out of 430,543 shared variants per sample, 83.7% of Ultima calls were shared with Illumina but this represented only 11.4% of Illumina high AF calls), whereas low AF variants showed 67.5% overlap (22,864 shared variants per sample) where only 0.88% overlapped with Illumina’s low AF variants, consistent with most low-frequency calls being platform-specific. Ultima exhibited 83,678 unique high AF variants per sample, compared with 3,336,667 for Illumina, indicating that Illumina is substantially more permissive in calling germline variation. Conversely, Illumina demonstrated 2,585,182 unique low AF variants per sample versus 11,004 for Ultima, consistent with a higher false positive rate (Figure 5B).

Low VAF variants were rare overall, with Ultima detecting an average of 337 per sample (Figure 5C). These calls were enriched for indel artifacts characteristic of the platform, highlighting a specific error mode. Notably, the proportion of indels classified as low VAF was 31.72% in Illumina, nearly twice that observed in Ultima (15.23%), indicating that Ultima is better at minimizing artifactual Indel calls compared to Illumina since in standard germline samples it is expected variants to be homozygous (VAF ∼ 100%) or heterozygous (VAF∼50%). Collectively, these results demonstrate that Ultima UG100 platform and pipeline achieves enhanced germline sensitivity while markedly reducing spurious low-frequency calls relative to Illumina’s.

Functional annotation of called SNPs did not reveal major differences between technologies: intronic variants represented the largest category (∼36%), followed by intergenic variants (∼32–33%), while protein-coding variants accounted for only ∼0.17–0.22% of calls in both platforms (Table 7), consistent with the distribution expected from whole-genome germline variant catalogs^21^.

#### Differential recovery of GWAS-catalogued IBD-associated variants across workflows

To evaluate clinical relevance, we curated a set of 89 germline variants located within 15 genes implicated in IBD, Crohn’s disease, and ulcerative colitis based on GWAS Catalog annotations (GRCh38 coordinates). These loci include variants in genes involved in immune regulation and intestinal barrier function such as L23R/IL12RB2, IL10, ZNF365, NKX2-3, NOD2, ATG16L1, PTPN2, TYK2, IL12B, TNFSF15, CARD9, NFKB1, and multiple HLA class II genes. Many of these loci were originally identified or confirmed in large GWAS and meta-analysis efforts in IBD research were used here as a predefined catalog of position of interest^22–25^. Five WGS samples were available for this comparative analysis (two Crohn’s disease and three ulcerative colitis samples).

Across the five samples, the number of catalogued variants detected per sample differed between workflows. On average, 18.4 variants from the curated set were detected per sample by Illumina (either Illumina-only or by both technologies). Of these, 16.8 were detected by Illumina only (i.e. not called by Ultima in that sample), and 1.6 on average were detected by both technologies. Ultima detected on average 1.6 variants per sample in this set, all of which were also detected by Illumina in the same sample; no variant was detected by Ultima only in any sample (Figure 5D and Table 8).

Thus, within this curated collection of IBD-associated loci, Illumina recovered a larger fraction of the 89 variants per sample than Ultima, with most detections being Illumina-only. This observation likely reflects differences in variant calling permissiveness and filtering thresholds between workflows, which may influence whether variants at specific loci pass germline calling criteria under the applied configuration. Importantly, this analysis evaluates detection within a predefined catalog rather than confirmed sample-specific truth variants; therefore, these results should be interpreted as a comparison of workflow behaviour at literature-reported position of interest, rather than a formal estimate of calling accuracy or sensitivity, which should be assessed using benchmarking datasets or validated genotype states of these loci in the analyzed samples.

## Discussion

Sequencing technologies are advancing through both the continued optimization of established platforms and the emergence of novel chemistries, collectively driving improvements in throughput, cost efficiency, and the diversity of approaches to data generation. This expansion directly enables AI-driven analytical frameworks to leverage large, high-fidelity omics data to extract disease-relevant biological signals. Realising this potential requires datasets of sufficient scale and clinical depth. Archival FFPE material underpins major multi-modal datasets including Owkin’s MOSAIC, as FFPE biobanks represent the largest source of clinically annotated, phenotypically diverse samples at the scale required for generalisable models. As datasets grow in number and scale alongside increasing platform diversity, rigorous head-to-head benchmarking of how sequencing chemistries and computational methods jointly determine biological capture becomes essential to deploy each technology appropriately and to define strategies for integrating data generated across them.

In this study, we generated a purpose-designed multi-modal dataset from human oncology and IBD FFPE specimens, spanning WTS, snRNA-seq, WES, and WGS, across both Ultima UG100 and Illumina NovaSeq platforms. We then performed a systematic head-to-head benchmarking of both platforms using clinically relevant workflows and platform-optimised analytical pipelines.

Illumina and Ultima differ fundamentally in their chemistries and error profiles. Illumina’s SBS chemistry performs stepwise nucleotide incorporation using 3′-blocked, dye-labeled nucleotides, producing highly accurate short reads with uniformly low indel rates, enabling reliable multi-mapping alignment and discrimination among homologous regions ^26,27^. By contrast, Ultima’s UG100 mnSBS platform employs non-terminating incorporation and optical endpoint scanning in an open-fluidics system, generating longer reads at lower per-base cost, but with characteristic insertion-dominant indels ^6,28^. These mechanistic distinctions are important; however, our data indicate that, in practice, their impact on biological inference is often modest and highly dependent on downstream analytical pipelines.

In bulk RNA-seq data from FFPE-derived human oncology and IBD specimens, Illumina demonstrated a slight over-representation of pseudogenes and certain non-coding transcripts, consistent with its ability to align multi-mapping reads confidently to homologous regions. Conversely, Ultima showed a modest enrichment for protein-coding gene assignment, presumably because its error profile reduces aligner confidence for ambiguous reads, pushing the mapping toward unique loci or discarding ambiguous alignments. Importantly, these platform-specific biases influenced only a small fraction of the transcriptome: the detection of lowly expressed genes, canonical biomarkers, oncogenic drivers, immune signaling genes, and pathway-level signals remained highly concordant. Nevertheless, we identified a group of genes totally absent from the Ultima dataset, some of which have been reported in the literature to have a role in disease biology or as biomarkers. As this group of genes did not display a significant content of long homopolymers, pseudoalignment methods (such as kallisto or salmon) could rescue some of these assignments as reported previously ^7^. This observation echoes previous cross-platform studies, which found slight differences in individual-gene detection, while the global expression patterns, known signatures and biological conclusions remained consistent. The preservation of relative expression levels across platforms indicates that the biological signal significantly outweighs any technology-specific noise, suggesting that data from Illumina and Ultima can be combined for meta-analysis without the use of traditional batch effect mitigation

For single nuclei transcriptome analysis, we assessed the performance of Ultima in a probe-based hybrid-capture assay, the Chromium SCR Flex, which targets pre-defined transcript regions instead of relying on full-length RNA. This approach greatly reduces dependence on RNA integrity and dramatically mitigates the impact of platform-specific sequencing error signatures. Consistent with prior studies that assessed non-fixed specimens, we found nearly-identical cell-type clustering, proportions, and UMI counts per cell, despite a ∼5-fold difference in raw read depth per cell ^29^. The extra depth on Ultima largely represented oversampling rather than additional unique transcripts. The only noted deviation occurred in a DLBCL cohort, where lymphoid subpopulations diverged modestly between platforms. Such differences may stem from the higher reads per cell, allowing for lymphocytes to fall below the qualification thresholds compared to cell types with a low total transcript count.

Our cross-modality assessment of WES and WGS highlights the technical equivalence of Ultima UG100 and Illumina for clinical genomic workflows, while uncovering distinct, pipeline-dependent performance profiles. In WES, both platforms achieved high-quality coverage and reliably captured all known clinically actionable oncogene hotspot mutations. However, Ultima UG100 exhibited a characteristic indel error bias toward lower-complexity regions, such as homopolymers, and employed a more conservative variant calling approach with DeepVariant, leading to fewer platform-specific unique calls compared to Illumina’s more permissive DRAGEN pipeline. Similarly, in WGS, UG100 demonstrated enhanced specificity for high-confidence germline variants while significantly reducing the burden of spurious low-frequency artifactual calls - showing a substantial reduction in potential false positives relative to Illumina. While Illumina maintains a sensitivity advantage for recovering a larger total fraction of documented GWAS-associated variants, the high precision and lower per-base cost of the Ultima platform provide a robust, scalable alternative for population-scale genomic studies, where minimizing false-positive noise is paramount.

Interpretations of the GIAB results should consider benchmarking context. GIAB reference is often considered a “gold standard,” but it was largely developed and validated using existing short-read sequencing technologies, primarily Illumina-based platforms. Because the GIAB truth sets are heavily influenced by the error profiles and biases of the technologies used to build them, they may not perfectly represent a “neutral” ground truth for a disruptive new chemistry like Ultima’s. In addition, Ultima’s pipeline may be at a disadvantage if it hasn’t been specifically optimized against the exact GIAB reference versions used in this study, particularly regarding its known sensitivity to homopolymer-related indel artifacts. Thus, a fair comparison would require orthogonal validation and cross-technology profiling of the same GIAB samples to uncouple chemistry-specific error modes from pipeline-dependent biases. It should also be noted that Ultima’s marginal lead in sensitivity in this specific benchmark likely reflects its higher raw sequencing depth in the GIAB dataset, as this advantage was not maintained in the subsequent equal-coverage cohort analysis.

It is notable that the UG100 platform evaluated here represents the first commercial implementation of mnSBS technology. Since completion of this study, Ultima has introduced the UG200 series, which incorporates updated amplification workflows, improved throughput, and refinements aimed at enhancing coverage uniformity and performance in challenging genomic regions. While our data provide a foundational benchmark for mnSBS chemistry in real-world FFPE applications, future evaluations will be required to determine the extent to which these hardware and workflow updates mitigate the indel and low-complexity sensitivities observed in UG100 and further close the precision gap in germline benchmarking. Importantly, the core biological concordance demonstrated here suggests that incremental chemistry and workflow refinements are likely to improve quantitative performance without altering overall biological fidelity.

For AI-scale multi-omic applications, the critical question is not whether platforms are identical at the base-call level, but whether they produce stable, interoperable biological representations that can co-exist in training and validation datasets with understood data analysis implications. Across modalities, our data indicate that UG100 captures disease clustering, pathway activation, immune composition and clinically actionable variants with high fidelity. Differences in background variant burden and ambiguous read assignment highlight the importance of harmonized bioinformatic pipelines and explicit calibration of variant-calling thresholds when integrating datasets across platforms. In large-scale modeling efforts, controlling systematic noise and maintaining reproducible signals may be as important as maximizing sensitivity.

This study has limitations. The cohort size is modest and restricted to FFPE specimens, limiting power to detect subtle differences in rare variant sensitivity or fine-grained cell-state annotation. Our use of platform-optimized pipelines reflects real-world practice but necessarily conflates chemistry and workflow effects. Broader validation across larger cohorts, fresh-frozen tissues, harmonized cross-processing strategies, and orthogonal truth datasets will be essential to fully characterize performance variability and optimize cross-platform integration.

Illumina remains the established benchmark for regulatory-grade sequencing and cross-study harmonization, supported by extensive validation and a mature analytical ecosystem. However, as sequencing demand shifts toward population-scale, AI-enabled analyses, cost-efficient and scalable alternatives are increasingly relevant. Our findings demonstrate that mnSBS-based sequencing, as implemented in UG100, achieves broad biological equivalence to established SBS platforms across transcriptomic and genomic applications. With the introduction of next-generation systems such as UG200, continued independent benchmarking will be critical to assess whether improvements in throughput, amplification chemistry, and coverage uniformity further narrow workflow-dependent performance gaps while preserving the biological fidelity demonstrated here.

Overall, these results position mnSBS technology as a viable and scalable contributor to the evolving multi-omic sequencing landscape, provided that platform-specific analytical considerations are transparently addressed and appropriately calibrated.

## Code availability

https://github.com/owkin/ultilumina-pilot/tree/main

## Acknowledgments

The authors would like to thank CeGaT and Ultima Genomics for providing sequencing services and technical support. These parties were involved solely in data generation and technical consultation; the study design, data analysis, and interpretation of results were conducted independently by the authors.

## Methods

### Data generation set up

#### Sample description and data flow

A comparative dataset was generated to enable a direct, head-to-head evaluation of sequencing performance between the Ultima UG100 and Illumina NovaSeq platforms. Fifteen samples representing five biological disease indications - diffuse large B-cell lymphoma (D), glioblastoma (G), muscle-invasive bladder cancer (M), ulcerative colitis (U), and Crohn’s disease (C) - were analyzed, with three donors per indication. Four sequencing modalities were included: single-nucleus RNA sequencing (snRNA-seq), whole transcriptome sequencing (WTS), whole exome sequencing (WES), and whole genome sequencing (WGS). WES was applied only to oncology samples, whereas WGS was performed on IBD specimens (U and C). To assess the technical reproducibility, one sample per sequencing modality was processed in duplicates.

All specimens were collected with informed consent by the local hospital biobank under institutional ethical approval (ethical approval 329_16B and 4607 from the FAU Ethical Board). They comprised FFPE blocks archived in the biobank for 1–5 years. Each block was sectioned within the local CRB laboratory. Two FFPE curls of 10 µm thickness were generated from each specimen, conserved at 4 °C in RNase-free conditions, and shipped to sequencing facilities at CeGaT (Tübingen, Germany) and/or Ultima Genomics (Fremont, CA, USA).

Library preparation for snRNA-seq, WTS, and WES was carried out at CeGaT (Tübingen, Germany). Libraries were sequenced on an Illumina NovaSeq system at CeGaT (NovaSeq 6000 for snRNA-seq and NovaSeq X Plus for bulk modalities). Identical aliquots of each library were then transferred to Ultima Genomics (Fremont, CA, USA), converted using Ultima-specific adapters and barcodes, and sequenced on the UG100 platform.

Genomic DNA and total RNA were isolated from FFPE tissue using the MagMAX™ FFPE DNA/RNA Ultra Kit (Thermo Fisher Scientific) according to the manufacturer’s instructions. Nucleic acid quantity was determined by the fluorescence-based assays, and integrity was assessed by fragment analysis using an Agilent Bioanalyzer or Fragment Analyzer system. For WGS, genomic DNA was extracted and evenly divided between the two facilities. One portion was processed and sequenced on the Illumina platform at CeGaT, and the other was independently processed and sequenced on the UG100 at Ultima Genomics.

#### Library preparation and sequencing settings

FFPE scrolls were used as the starting material for single-cell RNA sequencing. Cell concentration was determined using the Cellaca MX cell counter (Revvity). 8,000 cells per sample were loaded onto a Chromium chip and processed using the Next GEM Single Cell Fixed RNA Reagent Kits (10x Genomics) according to the manufacturer’s protocol. Libraries were sequenced on an Illumina NovaSeq 6000 platform using an S1 flow cell with paired-end reads (Read 1: 28 bp, Index 1: 10 bp, Index 2: 10 bp, Read 2: 90 bp).

Bulk transcriptome libraries were generated from 10 ng of total RNA using the SMART-Seq Stranded kit (Takara Bio), following the recommended workflow for cDNA synthesis, amplification, fragmentation, and indexing. For whole-exome sequencing, 50 ng of DNA was used as input for library construction with the Twist Human Core Exome + RefSeq + Mitochondrial Panel (Twist Bioscience), incorporating enzymatic fragmentation, adapter ligation, amplification, and target enrichment. Whole-genome libraries (for Illumina workflow) were prepared from 50 ng of DNA using the Twist Mechanical Fragmentation Library Preparation Kit (Twist Bioscience), including mechanical shearing, end repair, A-tailing, adapter ligation, and PCR amplification. Illumina bulk DNA libraries were sequenced on an Illumina NovaSeq X Plus platform using paired-end 2 × 101 bp reads. To ensure an accurate quality score estimation for the final base of each read, one additional cycle was intentionally added to both read 1 and read 2 relative to the standard 100-bp configuration. For the Ultima workflow, whole-genome libraries were prepared using the UG Solaris Flex DNA library preparation workflow, following the manufacturer’s recommendations. Nucleic acid quantity was determined by fluorescence-based assays, and integrity was assessed by fragment analysis. For each library, 10ng of genomic DNA was used as input. Indexed libraries were subsequently pooled to a final concentration of 750pM before bead preparation, QC and sequencing. Samples were sequenced using the UG 100 platform for 116 flow-cycles. Reads, prior to variant calling, were aligned CRAM files (Hg38), trimmed to remove redundant sequences, demultiplex and outputted as .CRAM files.

#### Ultima library conversion

Whole exome libraries were re-indexed with Ultima barcodes using the following protocol: UG Indexing PCR Library Preparation Protocol for Solaris Flex (https://cdn.sanity.io/files/l7780ks7/production-2024/0252074011f422caa89913e0d3d7a83f6c47c599.pdf). 10ng of the initial Twist Exome Library were amplified using indexing and universal primers (IDT Bioscience, PN10028413 and 10028399). Libraries were pooled into two pools at a final concentration of 750 pM, then seeded and clonally amplified on sequencing beads using a high-scale emulsion amplification tool. Samples were sequenced using the UG100 sequencing platform. Reads, prior to variant calling, were aligned CRAM files (Hg38), trimmed to remove redundant sequences, demultiplex and outputted as .CRAM files for variant calling.

### Bulk RNA sequencing

#### Ultima read downsampling strategy

To enable fair comparison between Ultima and Illumina sequencing technologies, a systematic downsampling approach was implemented to normalize read counts across platforms. Ultima sequencing data was initially converted from CRAM format to FASTQ using SAMtools version 1.22 with the command samtools fastq --reference to ensure proper reference-based decompression. The downsampling strategy employed Seqtk to randomly subsample Ultima reads to match the corresponding Illumina sample read counts, as determined from MultiQC post-alignment reports. For each Ultima sample, the target read count was established by identifying the corresponding Illumina sample. The downsampling process utilized a fixed random seed to ensure reproducibility and was performed using the command seqtk sample -s100 with parallel processing capabilities. Quality control validation was performed by comparing read counts between original CRAM files and converted FASTQ files using Samtools view count functionality, ensuring data integrity throughout the conversion and downsampling pipeline. This approach ensured that both sequencing technologies were analyzed with equivalent sequencing depth, eliminating potential bias due to different library sizes while maintaining the biological signal necessary for comparative analysis.

#### Quality control and adapter trimming

Raw FASTQ files underwent comprehensive quality assessment using FastQC version 0.12.1 to evaluate multiple quality metrics including per-base sequence quality, per-sequence quality scores, sequence length distribution, sequence duplication levels, overrepresented sequences, adapter contamination, and GC content distribution. The quality control analysis was performed on both paired-end and single-end reads with automatic detection of sequencing configuration. Quality control reports were generated in HTML format for individual samples and subsequently aggregated using MultiQC version 1.17 to provide a comprehensive overview of pre-alignment quality metrics across all samples in the dataset.

#### Read alignment

Read alignment was performed using STAR version 2.7.9a with the human reference genome GRCh38 and GENCODE version 39 annotation. The STAR index was constructed with a splice junction database overhang of 100 base pairs using the reference genome assembly and annotation files. The alignment process was configured to generate unsorted BAM files with gene count quantification enabled using the --outSAMtype BAM Unsorted --quantMode GeneCounts parameters. Additional alignment parameters included a unique mapping quality score threshold (MAPQ) of 60 (--outSAMmapqUnique 60) to ensure high-confidence alignments.

The alignment process incorporated specialized parameters for chimeric alignment detection to support fusion gene analysis, including a minimum chimeric segment length of 10 base pairs (--chimSegmentMin 10), chimeric output type within BAM files with soft clipping (--chimOutType WithinBAM SoftClip), and junction overhang parameters optimized for fusion detection. The alignment utilized the GENCODE v39 annotation file for splice junction database construction and employed sophisticated scoring parameters for chimeric junction detection including minimum chimeric scores, maximum score drops, and junction-specific scoring criteria. The alignment process automatically detected paired-end versus single-end sequencing data and applied appropriate alignment strategies for each configuration.

#### Post-alignment quality control

Comprehensive post-alignment quality assessment was performed using RSeQC (RNA-seq Quality Control) version 5.0.4 to evaluate multiple aspects of alignment quality and library characteristics. The RSeQC analysis included junction saturation analysis to assess the completeness of splice junction detection, read distribution analysis to evaluate the proportion of reads mapping to different genomic features (exons, introns, intergenic regions, and untranslated regions), inner distance analysis to assess fragment size distribution and library preparation quality, and read duplication analysis to identify potential PCR artifacts or over-amplification issues.

Additional quality control metrics were generated using SAMtools version 1.21 for chromosome-level alignment statistics and BAM file indexing. The post-alignment quality control process incorporated strand-specific analysis using RSeQC’s infer experiment functionality to determine library strand orientation and validate the expected strandedness of the sequencing libraries. All quality control metrics were aggregated using MultiQC version 1.17 to generate comprehensive post-alignment quality reports with standardized visualizations and summary statistics.

In addition, quantification on the percentage of total reads assigned to different categories was conducted from the BAM QC output. Specifically, the proportion of unique reads (uniquely_mapped), multimapped (multimapped) and unmapped reads (unmapped_other + unmapped_tooshort + multimapped_toomany) normalized by total number of reads, high quality primary alignment (mapq_gte_mapq_cut_unique) normalized by total records and percentage of spliced/non spliced reads (splice_reads/non.splice_reads) normalized by mapq_gte_mapq_cut_unique.

#### Count matrix generation

Raw gene count matrices were generated from STAR alignment output by extracting gene-level quantification data from the ReadsPerGene.out.tab files produced during the alignment process. The count extraction process utilized the gene count quantification mode enabled during STAR alignment, which provided counts for each gene based on the GENCODE v39 annotation. The count matrix generation process aggregated individual sample count files into a comprehensive matrix format with genes as rows and samples as columns, ensuring consistent gene identifier mapping across all samples and technologies.

The count matrix generation incorporated strand-specific information derived from RSeQC infer experiment analysis to properly handle stranded sequencing libraries and ensure accurate gene count assignment. The process utilized Python-based scripts with pandas for data manipulation and integration, ensuring robust handling of large-scale count data and proper gene identifier mapping. The final raw count matrix was generated in tab-separated value format with Ensembl gene identifiers as row names and sample identifiers as column headers, providing a standardized format suitable for downstream differential expression analysis and statistical modeling.

#### Batch effect correction and normalization

Batch effect correction was performed using Combat-Seq algorithm, implemented through the inmoose.pycombat package. The batch correction process identified sequencing technology (Illumina vs Ultima) as the primary batch variable, while preserving biological variation associated with indication groups (C, D, G, M, U) as covariates in the correction model. The CombatSeq algorithm was applied to merged raw count matrices containing both Illumina and Ultima samples, using the technology column as the batch variable and indication as the covariate of interest. The correction process generated batch-corrected count matrices that were subsequently subjected to VST normalization using pydeseq2 (version 0.4.0) to ensure compatibility with downstream statistical analyses. The batch correction workflow included both corrected and uncorrected versions of normalized matrices to enable comparative assessment of the impact of batch correction on downstream analyses, including differential expression analysis and distance-based comparisons.

#### Calculating distance between pairs for biological signal comparison

Distance-based analysis was implemented to assess the accuracy of biological signal preservation between sequencing technologies across different indications. Euclidean distances were calculated in both gene expression space and principal component analysis (PCA) space to evaluate within-indication sample similarity. The analysis employed two complementary approaches: pairwise distance calculation in the original gene expression space using all genes with non-zero variance, and distance calculation in reduced-dimensional PCA space using the first 10 principal components. For each indication group (C, D, G, M, U), all possible pairwise combinations of samples within the same technology were identified, and Euclidean distances were computed using the formula d = √(Σ(xi - yi)²) where xi and yi represent the expression vectors for samples i and j. The distance calculations were performed separately for each sequencing technology to enable direct comparison of biological signal consistency. Additionally, the analysis included merged datasets with and without batch correction to assess the impact of batch correction on biological signal preservation.

#### Differential gene expression analysis

Differential expression analysis was performed using DESeq2 (version = 0.4.0), to identify genes significantly differentially expressed between indication groups within each sequencing technology. The analysis employed a negative binomial model with Wald test statistics to assess differential expression, using the DESeqDataSet class for data preparation and model fitting. For each technology (Illumina and Ultima), separate analyses were conducted comparing each indication against all other indications (e.g., C vs others, D vs others, G vs others, M vs others, U vs others). The analysis design matrix included indication as the primary factor of interest, with samples grouped into binary categories (target indication vs all others). Genes with zero counts across all samples were automatically filtered out prior to analysis. Statistical significance was determined using adjusted p-values (padj < 0.05) with Benjamini-Hochberg correction for multiple testing. Results were categorized into significant and non-significant gene sets, with additional classification of upregulated and downregulated genes based on log2 fold change direction.

We also ran differential expression analysis using technology as the primary factor of interest, correcting for subject ID.

### Single nuclei RNA seq

#### Sequencing processing and quality control

Raw sequencing data were processed using Cell Ranger (version 9.0.1). FASTQ files were generated from the raw BCL files and aligned to the reference genome. Cell Ranger performed barcode assignment, UMI deduplication, and gene quantification, producing both raw and filtered gene-expression matrices for downstream analysis. These matrices were loaded into scANVI 1.11.2 ^30^. We performed quality control (QC) for each sample in each indication separately to account for differences in data quality across the different samples. The analysis was performed according to the recommendations described in Heumos, Schaar et. al ^31^.

We apply QC filters to both datasets. First, QC metrics for genes and cells are calculated and subsequently, automatic QC based on mean absolute deviations (MAD) thresholding was performed. Cells are filtered using a MAD threshold of 5 for log1p total counts, log1p n genes per count and the percentage of counts in the top 20% expressed genes. Additionally, we filter cells which exceed 8% mitochondrial counts and cells which are flagged as mitochondrial outliers using a 3 MAD threshold. Doublet are detected using Scrublet 0.2.3 and are filtered using a doublet probability threshold of 0.2. All filtered samples are then merged into a single AnnData object for subsequent analysis. Next, we calculated for the merged AnnData object, a k nearest neighbor graph using 15 neighbors on the 50 dimensional PCA embedding serving as input to obtain a UMAP embedding of the entire dataset containing both samples measured with Ultima and Illumina. No subsequent batch correction was performed.

Annotation of the single-nuclei data was done on all cells after combining all the data across indications, patients and technologies in order to gain the highest resolution by having the largest number of cells. This was done before applying QC filters and removal of doublets. Consecutive clustering method was used where clusters from different subset of cells and resolutions were assigned into cell types based on canonical marker genes. Some clusters were identified as doublets via manual annotation, and removed from further analysis after applying the QC procedure.

#### Clustering and annotations

For cell-type annotation, we applied scANVI, using manually curated reference annotations. An established four-level hierarchical ontology was used to assign cell identities at multiple resolutions, with primary annotations generated at the fine-grained level and coarser levels inferred from the hierarchy. Cells flagged as potential doublets by manual review were retained in the processed data but annotated accordingly. The resulting corrected, filtered, and normalized dataset was used for all subsequent analyses.

In order to assess whether both technologies capture the same distribution of detected cell types in each sample, we performed two one-sided t-test to test for non-independence of the two technologies with a p-value threshold of 0.05 using statsmodels version 0.14.4 on a uniform random sample with a bootstrapping sample size of 500 for each indication and dataset. Next, we inspected cell types which are flagged as being from different distributions to assess the absolute differences in abundance in both technologies and analyse differences in their gene expression distribution across annotation levels 2, 3 and 4.

### Whole Exome/Genome data pre-processing

#### Ultima preprocessing

All data were automatically aligned on the instrument server using Ultima Aligner to Hg38 (https://github.com/Ultimagen/healthomics-workflows/blob/main/workflows/trim_align_sort/howto-ua-align-sort.md). WGS data analysis was performed using Ultima adjusted EfficientDeepVariant with a FFPE dedicated model (s3://genomics-pipeline-concordanz-us-east-1/deepvariant/model/somatic/wgs/ffpe/deepvariant-ultima-somatic-wgs-ffpe-model-v1.3.ckpt-890000.onnx) and an un-matched normal sample to filter as many germline SNV/indel as possible. WES data analysis was performed using Ultima adjusted EfficientDeepVariant with a dedicated germline model (s3://genomics-pipeline-concordanz-us-east-1/deepvariant/model/germline/v1.3/model.ckpt-890 000.dyn_1500.onnx).

The DeepVariant analysis utilized the GRCh38 reference genome and generated VCF files containing variant calls with comprehensive quality metrics and annotations.

#### Illumina processing

Illumina whole exome sequencing data underwent comprehensive preprocessing using DRAGEN version 07.031.676, a high-performance bioinformatics platform optimized for accelerated genomic analysis. The DRAGEN pipeline processed paired-end FASTQ files through an integrated workflow encompassing read alignment, duplicate marking, variant calling, and quality control analysis. The preprocessing utilized the GRCh38 reference genome with alternative contigs and employed a graph-based alignment approach with hash table optimization for enhanced mapping accuracy and computational efficiency.

The DRAGEN preprocessing pipeline incorporated multiple quality control and filtering steps including duplicate marking, mapping quality assessment, and target region analysis with 30-base pair padding around exonic regions. The variant calling component utilized DRAGEN’s somatic variant caller with systematic noise filtering using a panel of normals, germline tagging capabilities, and tumor-only analysis mode (only for Whole Exome Sequencing data). Quality filtering parameters included minimum allele frequency thresholds, base quality requirements, mapping quality thresholds, and read support requirements (minimum 3 supporting reads).

The pre-processing generated comprehensive quality metrics including mapping statistics, coverage analysis and variant calling summaries. Additional quality control included transition/transversion ratio analysis, heterozygosity assessment, and chromosome-specific variant distribution analysis. The pipeline also incorporated structural variant calling, copy number variation analysis, and microsatellite instability assessment, with all outputs formatted as VCF files containing standardized FORMAT fields for allele depth (AD), allele frequency (AF), read depth (DP), and strand bias information (SB) for downstream analysis and interpretation.

#### GC content analysis

GC bias was quantified from aligned BAM/CRAM files using Picard CollectGcBiasMetrics with a GRCh38 reference FASTA (Homo_sapiens_assembly38.fasta). We computed windowed GC content across the reference and reports, for each 1% GC bin, the number of windows, read starts, and normalized coverage (coverage normalized to the genome-wide mean), together with summary dropout metrics (AT_DROPOUT and GC_DROPOUT). For interpretability, we additionally aggregated 1% bins into low (<30%), mid (30–70%), and high (>70%) GC strata and summarized per-sample and per-platform read-start representation and mean normalized coverage in these strata. Between-platform differences in dropout metrics were assessed using two-sided Mann–Whitney U tests.

#### Functional annotation

We benchmarked Illumina and Ultima whole-exome variant callsets using VEP-annotated VCFs containing the INFO/CSQ field (Ensembl Variant Effect Predictor). Variants were parsed by streaming bgzipped VCFs; for each record we used the first alternate allele and the first CSQ transcript entry, with CSQ subfields mapped using the header-provided ’Format’ specification. Functional categories (protein-coding, missense, nonsense, splice-site, synonymous, frameshift, inframe indel, UTR, intronic, intergenic) were derived from VEP Consequence terms using a standard priority ordering, and summarized per sample. We evaluated two analysis modes: (i) QC-only, including variants with FILTER=PASS or ’.’, and (ii) QC+HIGH/MODERATE, additionally restricting to variants with VEP IMPACT in {HIGH, MODERATE}. Sample pairing across technologies used harmonized base identifiers derived from file stems (removing Illumina run suffixes and mapping ER_G_003 curl2 to a replicate key), and overlap was computed on exact (CHROM, POS, REF, ALT) matches. Ti/Tv was computed on SNVs as the ratio of transitions (A↔G, C↔T) to transversions, with indels/MNPs tracked separately.

### Whole Exome Sequencing Downstream analysis

#### Variant preprocessing and quality filtering

Raw VCF files from both Illumina DRAGEN and Ultima DeepVariant pipelines underwent systematic preprocessing to ensure consistent variant calling standards and remove low-quality variants. For Illumina data, variants were filtered to retain only those with FILTER status “PASS” and variant allele frequency (VAF) ≥ 0.1, as determined from the AF field in the FORMAT column. Ultima variants underwent identical filtering criteria, with VAF extracted from the VAF field in the FORMAT column. The preprocessing step resulted in high-confidence variant calls suitable for downstream comparative analysis.

#### Variant annotation and functional classification

Pre-processed variants underwent comprehensive functional annotation using the Ensembl Variant Effect Predictor (VEP) version 108 with the GRCh38 reference genome. VEP annotation included gene symbols, functional consequences, impact predictions, biotype information, and protein-level changes.

#### Germline variant filtering

Germline variants were identified and filtered using technology-specific criteria to focus analysis on somatic mutations. For Illumina data, variants were classified as germline if they contained “GermlineStatus=Germline_DB” in the INFO field, indicating database-confirmed germline status. For Ultima data, variants were classified as germline if their ID field contained rs identifiers (rs), representing known population variants.

#### Tumor suppressor gene and oncogene classification

Variants were classified into functional categories using curated gene lists and hotspot mutation databases. Tumor suppressor genes (TSGs) were identified using the IntOGen database, including well-established cancer genes such as TP53, BRCA1, BRCA2, and PTEN. Oncogene hotspot mutations were identified using a comprehensive database of 1,607 known cancer hotspot positions across 426 genes ^32–34^, including specific protein changes associated with oncogenic activity. Variants were categorized as “TSG” if they occurred in tumor suppressor genes, “Oncogene” if they matched known hotspot positions and protein changes, or “Other” for variants not falling into these categories. This classification enabled focused analysis on clinically relevant mutation types.

#### Comparative variant analysis

Variant overlap analysis was performed to compare mutation detection between Illumina and Ultima technologies. For each sample, variants were grouped by functional category (TSG, Oncogene, Other) and technology-specific detection was assessed.

#### Manual variant inspection and validation

A subset of hostpot mutations was selected for manual inspection using the Integrative Genomics Viewer (IGV) to validate variant calling accuracy and assess potential false positives or false negatives. IGV visualization enabled assessment of read alignment quality, variant allele frequency, and potential sequencing artifacts. Manual inspection results were integrated into the final variant analysis to ensure high-confidence variant calls for downstream clinical interpretation.

### Whole Genome Sequencing Downstream analysis

#### Variant annotation and quality control

Raw VCF files from both Illumina DRAGEN and Ultima DeepVariant pipelines underwent systematic annotation and quality control to ensure consistent germline variant calling standards. Variants were first filtered to retain only those with FILTER status “PASS” to remove low-quality calls and reduce computational overhead for downstream processing. The PASS filtering step was performed using bcftools version 1.22. Following quality filtering, variants underwent comprehensive functional annotation using the Ensembl Variant Effect Predictor (VEP) version 108 with the GRCh38 reference genome and offline cache, enabling consistent annotation across both sequencing technologies.

#### Population frequency annotation and germline variant identification

VEP-annotated variants were further annotated with population allele frequencies using gnomAD v4.0 to enable germline variant identification and filtering. The gnomAD annotation was performed using bcftools annotate with exact pair-logic matching, adding AF and AF_grpmax (maximum AF across human populations) fields to the INFO column. This annotation step ensured consistent population frequency data across both Illumina and Ultima technologies, enabling standardized germline variant filtering based on population allele frequencies.

#### Variant categorization and technology comparison

Variants were systematically categorized based on variant allele frequency (VAF) and population frequency thresholds to enable comparative analysis between sequencing technologies. Variants were classified into four categories: “low_VAF” for variants with VAF < 0.1, “high_AF” for variants with gnomAD AF_grpmax > 0.01, and “low_AF” for variants with gnomAD AF_grpmax ≤ 0.01. This categorization enabled direct comparison of variant detection patterns between Illumina and Ultima technologies across different variant classes.

#### Sensitivity and specificity assessment through intersection analysis

Sensitivity and specificity of variant detection were assessed through comprehensive intersection analysis comparing variant calls between Illumina and Ultima technologies across all variant categories, performed on a per-sample basis. For each sample, variants were grouped by technology-category combinations (e.g., Illumina_low_VAF, Ultima_high_AF) and intersection analysis was performed to quantify shared variants, technology-specific variants, and cross-category matches. This approach enabled calculation of sensitivity metrics by identifying variants detected by both technologies in the high_AF category (supposed true positives), variants detected by only one technology in the high_AF category (false negatives for the missing technology). Specificity was assessed by analyzing the distribution of low_VAF variants (considered false positives) and their detection patterns, with cross-ensemble matches (e.g., Illumina_low_VAF ∩ Ultima_high_AF) providing insights into technology-specific bias patterns where one technology calls low-quality variants while the other correctly identifies high-quality variants.

#### Clinical relevance with regard to IBD GWAS ground truth

To benchmark germline variant detection between the two sequencing technologies on the same DNA samples, we defined a ground-truth set of expected IBD-associated variants. We started from a curated list of genes of interest implicated in inflammatory bowel disease (IBD) and related pathways (innate and adaptive immunity, autophagy, cytokine signalling, antigen presentation). The list comprised 15 genes: NOD2, IL23R, ATG16L1, IRGM, CARD9, TNFSF15, PTPN2, IL10, HLA-DRB1, HLA-DQA1, IL12B, TYK2, NKX2-3, ZNF365, and SLC39A8. We then used the GWAS Catalog (NHGRI-EBI Catalog of published genome-wide association studies; https://www.ebi.ac.uk/gwas/; Sollis et al., Nucleic Acids Res. 2023) to retrieve reported associations for three EFO traits: inflammatory bowel disease (EFO_0003767), Crohn disease (EFO_0000384), and ulcerative colitis (EFO_0000725). Restricting to associations whose reported or mapped gene was in our gene set. This yielded a list of variant identifiers (rs IDs) and genomic positions (GRCh38). We resolved reference and alternate alleles (REF/ALT) for each rs ID via NCBI dbSNP and built a final variant set as unique identifiers. The ground-truth set contained 89 IBD-associated germline variants.

We used whole-genome sequencing (WGS) data from IBD patients with the same samples replicated in both technologies. We analyzed five samples: two with Crohn disease and three with ulcerative colitis.

For each sample, we matched the ground-truth variant set against technology-specific summary tables (Illumina and Ultima) using the predefined variant identifier. We classified each ground-truth variant in each sample as detected by Illumina only, Ultima only, or by both. This yields, per sample, the counts of variants detected only by Illumina, only by Ultima, and by both, which we then used to assess the relative sensitivity: variants detected by both technologies provided a consensus set; variants detected by only one technology indicate technology-specific detection. We also produced a variant-level table listing each detected variant with sample ID and technology (Illumina, Ultima, or Both) for downstream analysis.

#### Indel benchmarking against GIAB

We compared indel calls from each platform against the GIAB reference set for HG001 (NA12878) on GRCh38 (NIST v3.3.2)^20^. To ensure a fair comparison and avoid bias from platform-specific filtering, we used the raw, unfiltered per-sample VCF files that serve as input to our VEP and gnomAD annotation pipelines, rather than PASS-filtered or post-processed summary tables. For each platform we concatenated indels from all per-sample VCFs and deduplicated by unique variants (so each normalized indel was counted once). Variants were normalized before matching: REF/ALT were left-aligned and trimmed to a minimal representation following standard VCF normalization. Matching was performed on a unique key (CHROM_POS_REF_ALT) derived from the normalized variant. For each technology we computed true positives (TP), false positives (FP), and false negatives (FN) against the 571,515 GIAB indels, and derived sensitivity (TP/(TP+FN)), precision (TP/(TP+FP)), and F1-score.

---

**Supplementary Figure 1.**
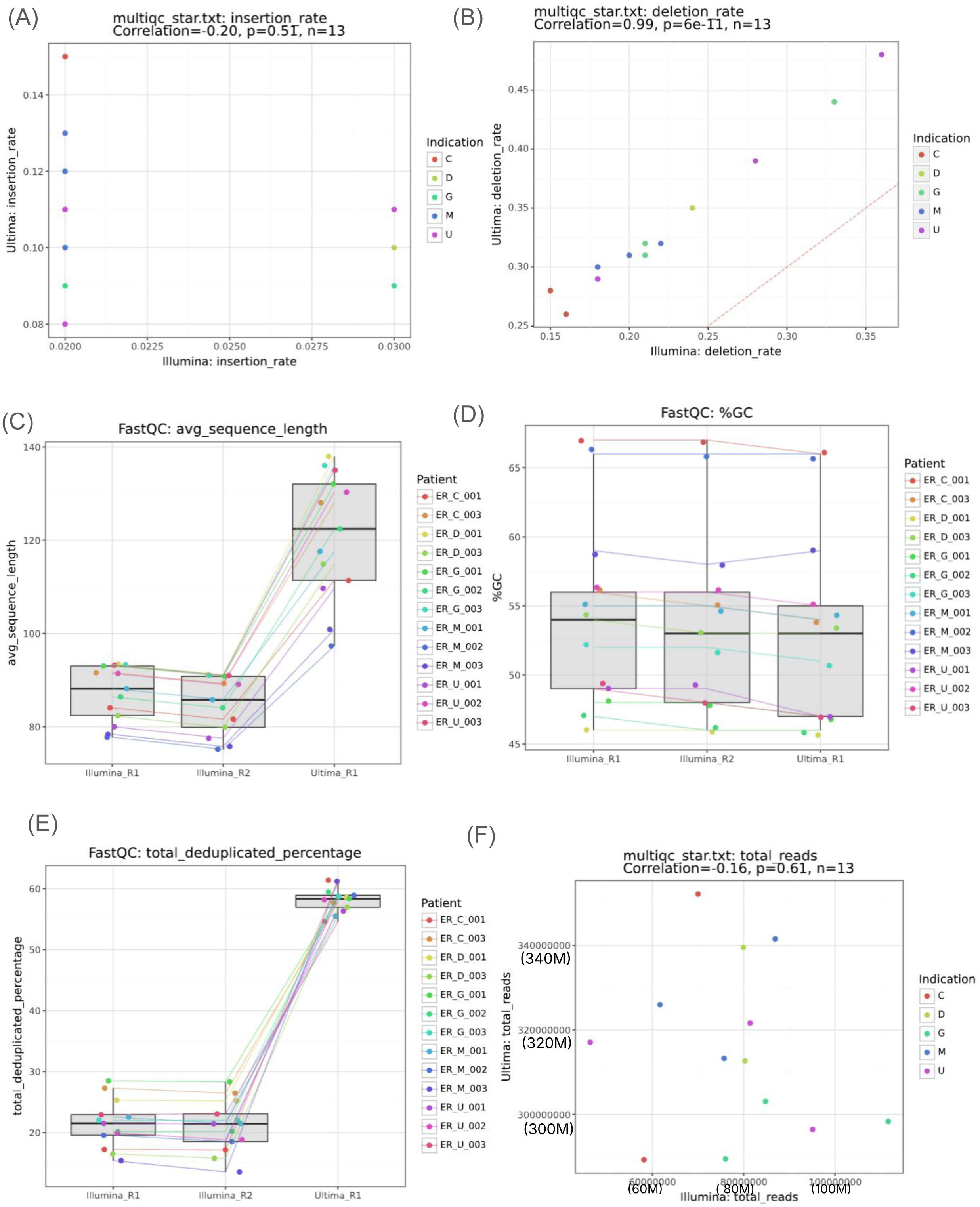
Bulk RNA-seq. QC sequencing bulk RNA-seq **(A)** Insertion rates **(B)** deletion rates. Red dotted line shows similar values, **(C)** average sequence length **(D)** Percentage GC **(E)** Percentage of deduplicated reads **(F)** Total number of reads Colors by disease indication (A, B, F) or patient (C, D, E).

**Supplementary Figure 2.**
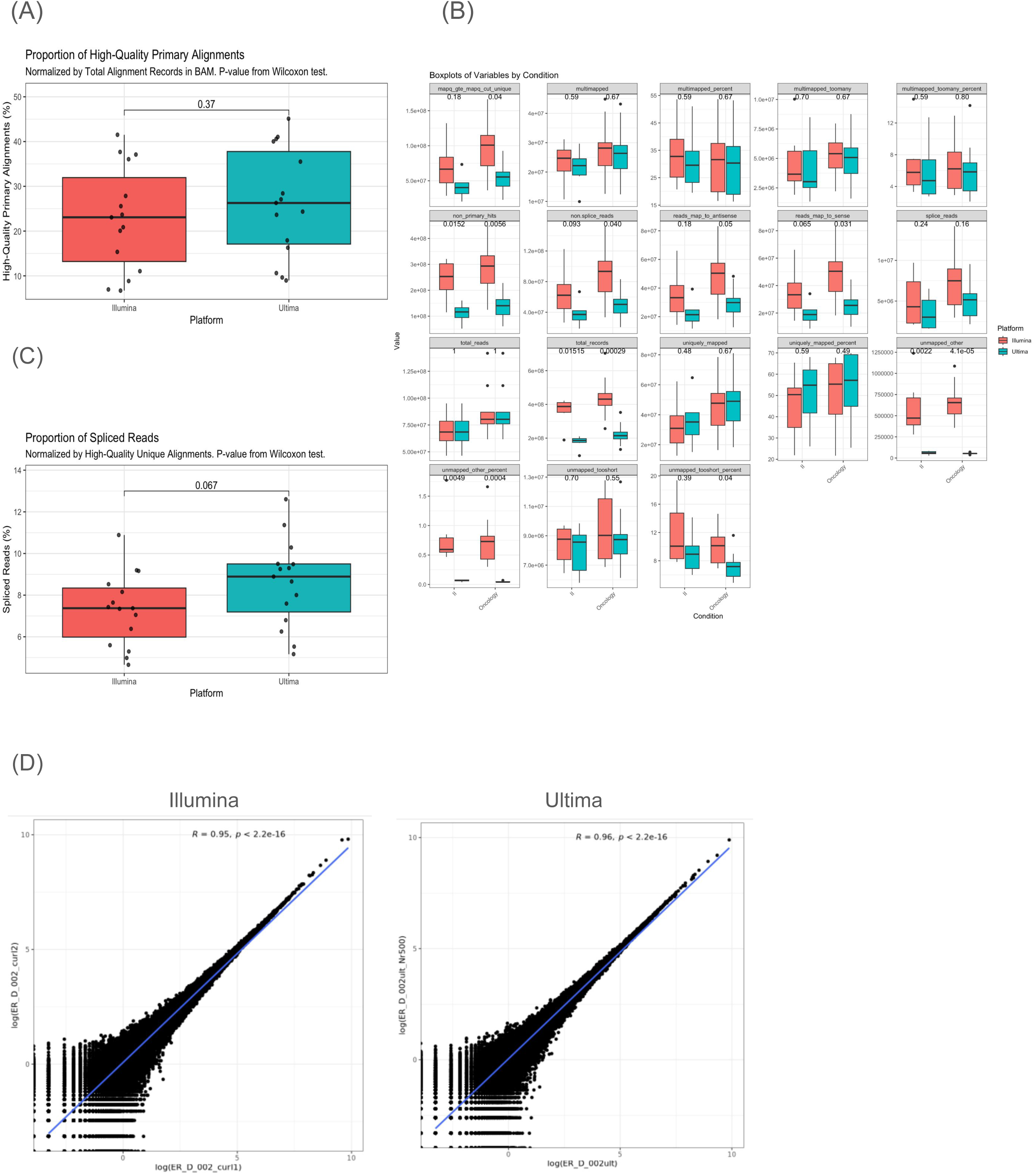
Bulk RNA-seq. **(A)** Proportion of high quality reads **(B)** Number of reads across QC categories for each platform, stratified by main disease group **(C)** Proportion of spliced reads **(D)** Gene expression counts from technical replicates

**Supplementary Figure 3.**
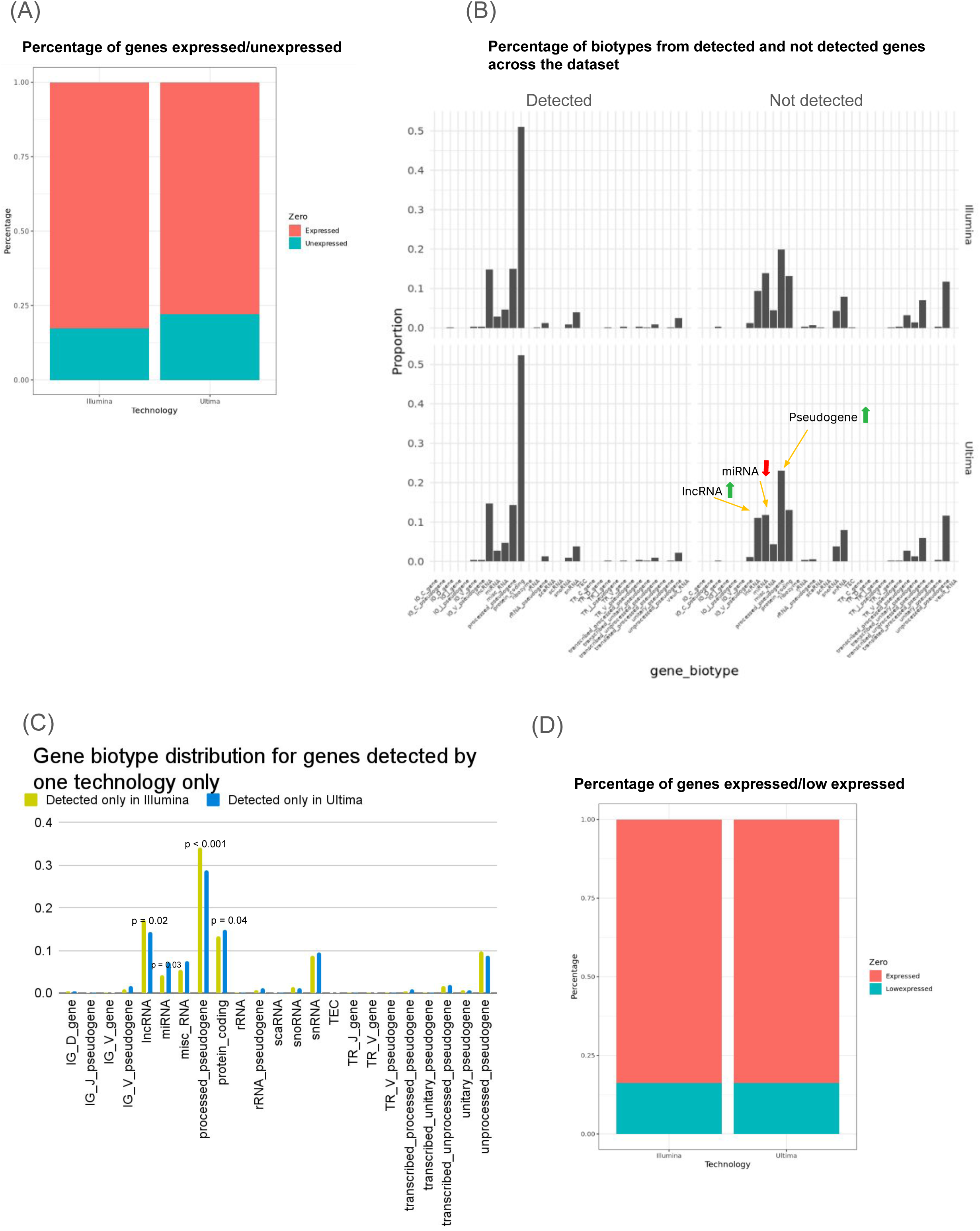
Bulk RNA-seq. **(A)** Proportion of expressed and unexpressed genes in each technology **(B)** Biotype distribution across expressed and unexpressed genes **(C)** Proportion of gene biotypes among genes detected in one technology only **(D)** Proportion of expressed and low expressed (< 10 counts) genes.

**Supplementary Figure 4.**
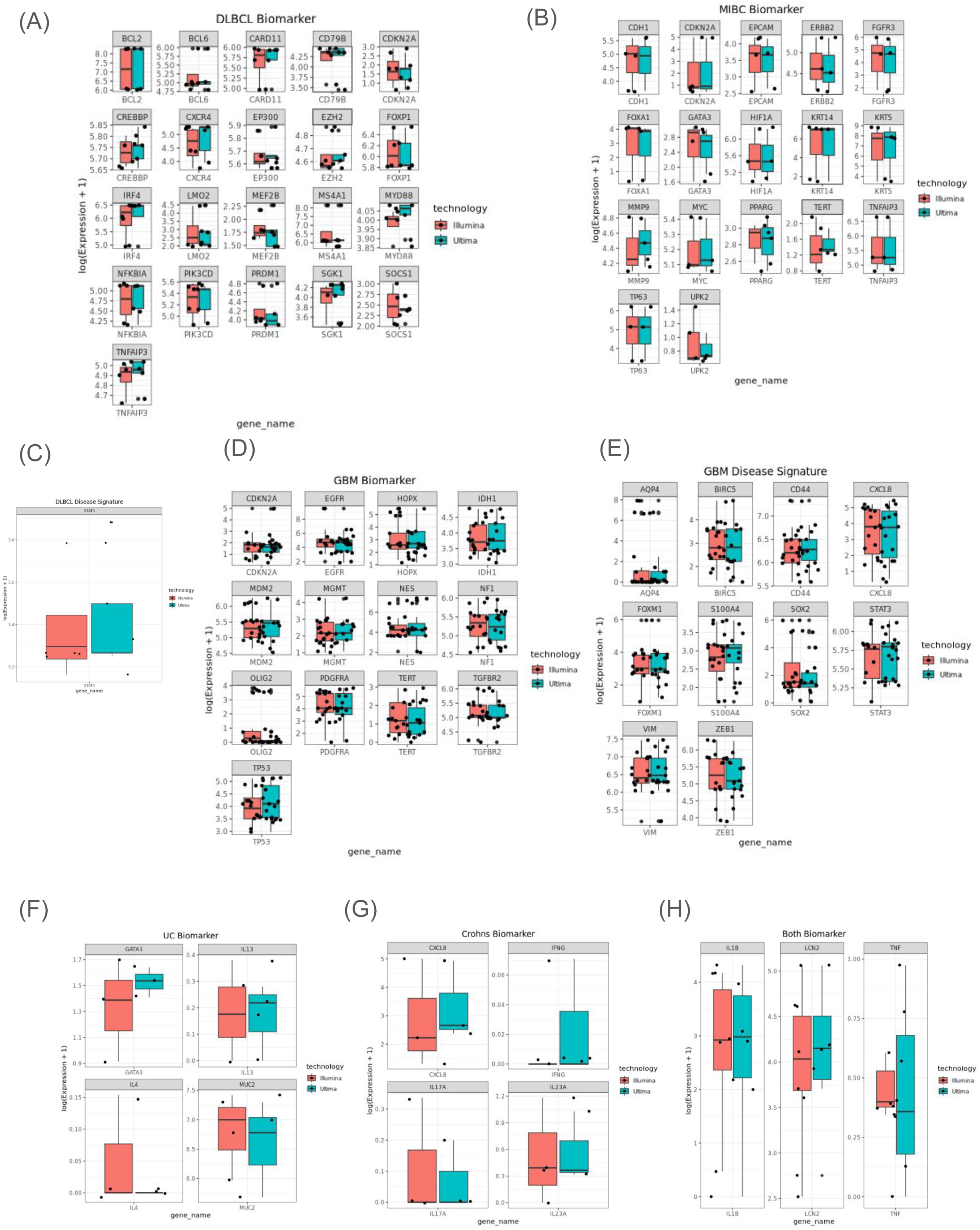
Bulk RNA-seq. Gene expression levels across selected genes of interest for DLBCL (**A** and **B**), GBM (**C** and **D**), UC (**E**), C (**F**) and IBD (**G**)

**Supplementary Figure 5.**
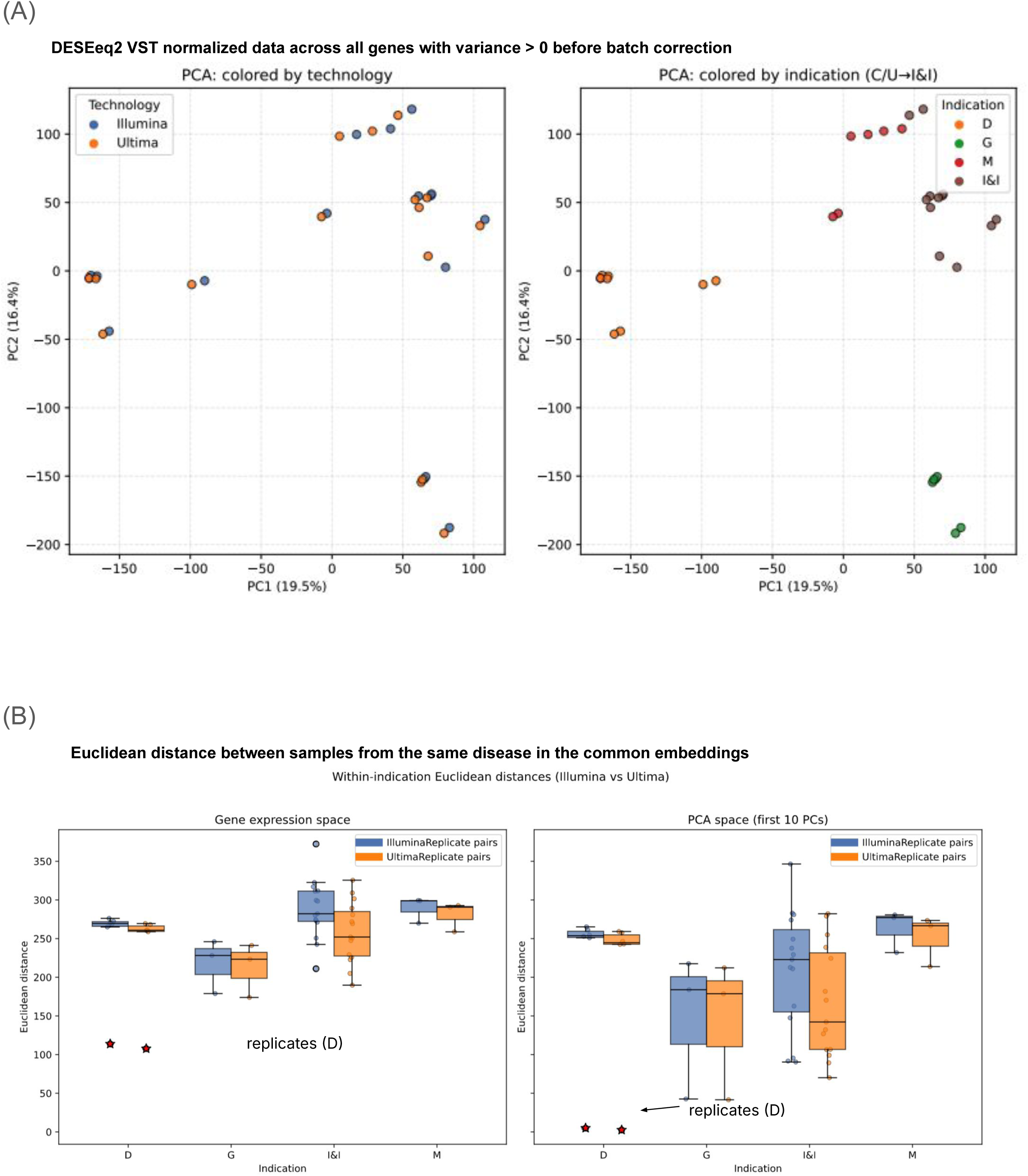
Bulk RNA-seq. **(A)** PCA from normalized gene expression profiles from all samples, colored by technology (left) and indication (right) **(B)** Euclidean distance in gene expression space (left) and PCA space (right) across from all samples. Replicates are represented by a red star.

**Supplementary Figure 6.**
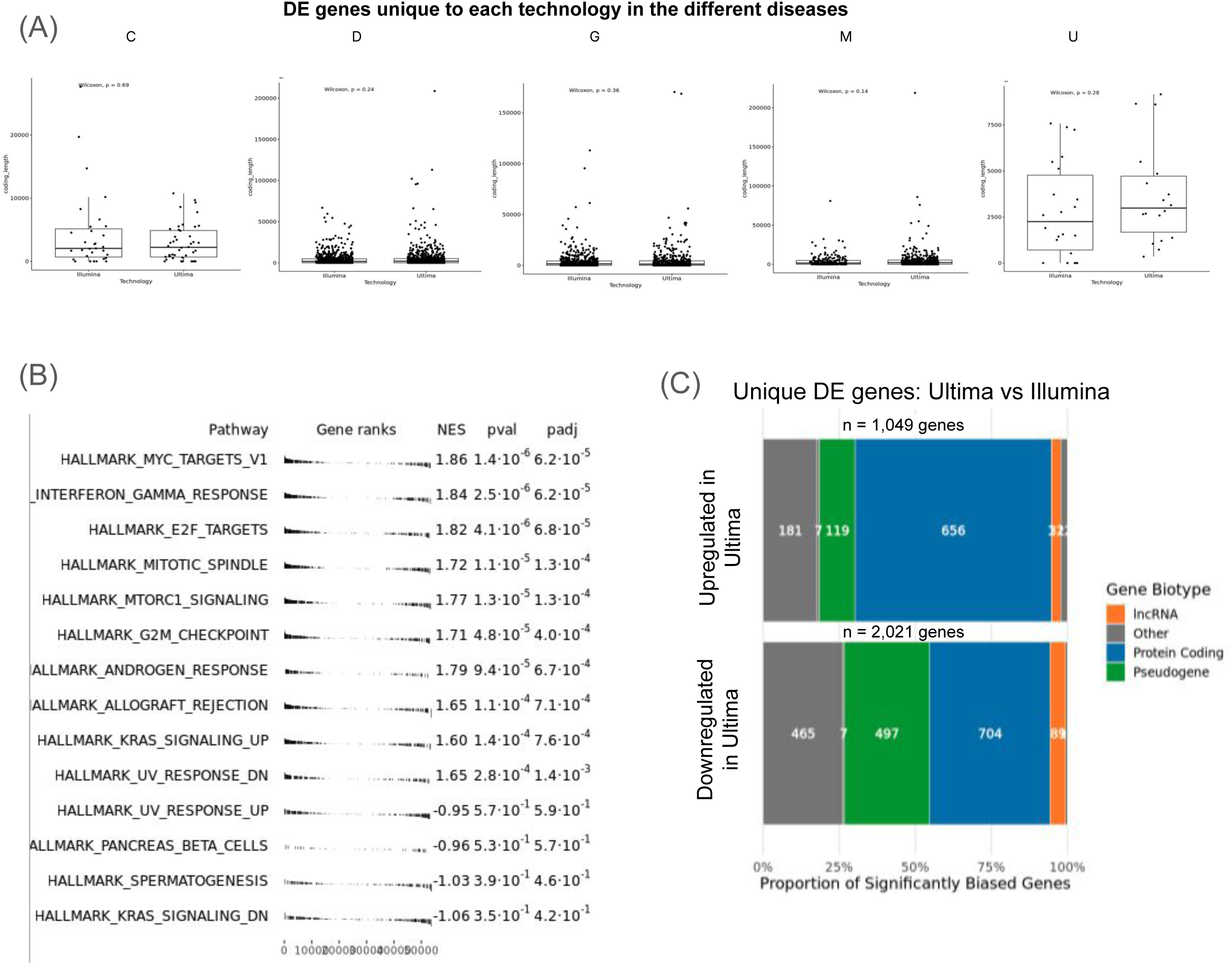
Bulk RNA-seq. **(A)** Distribution of gene length across diseases for each technology **(B)** Gene set enrichment analysis from differential expressed genes between Ultima and Illumina **C)** Biotype distribution of genes differentially expressed between Ultima and Illumina.

**Supplementary Figure 7.**
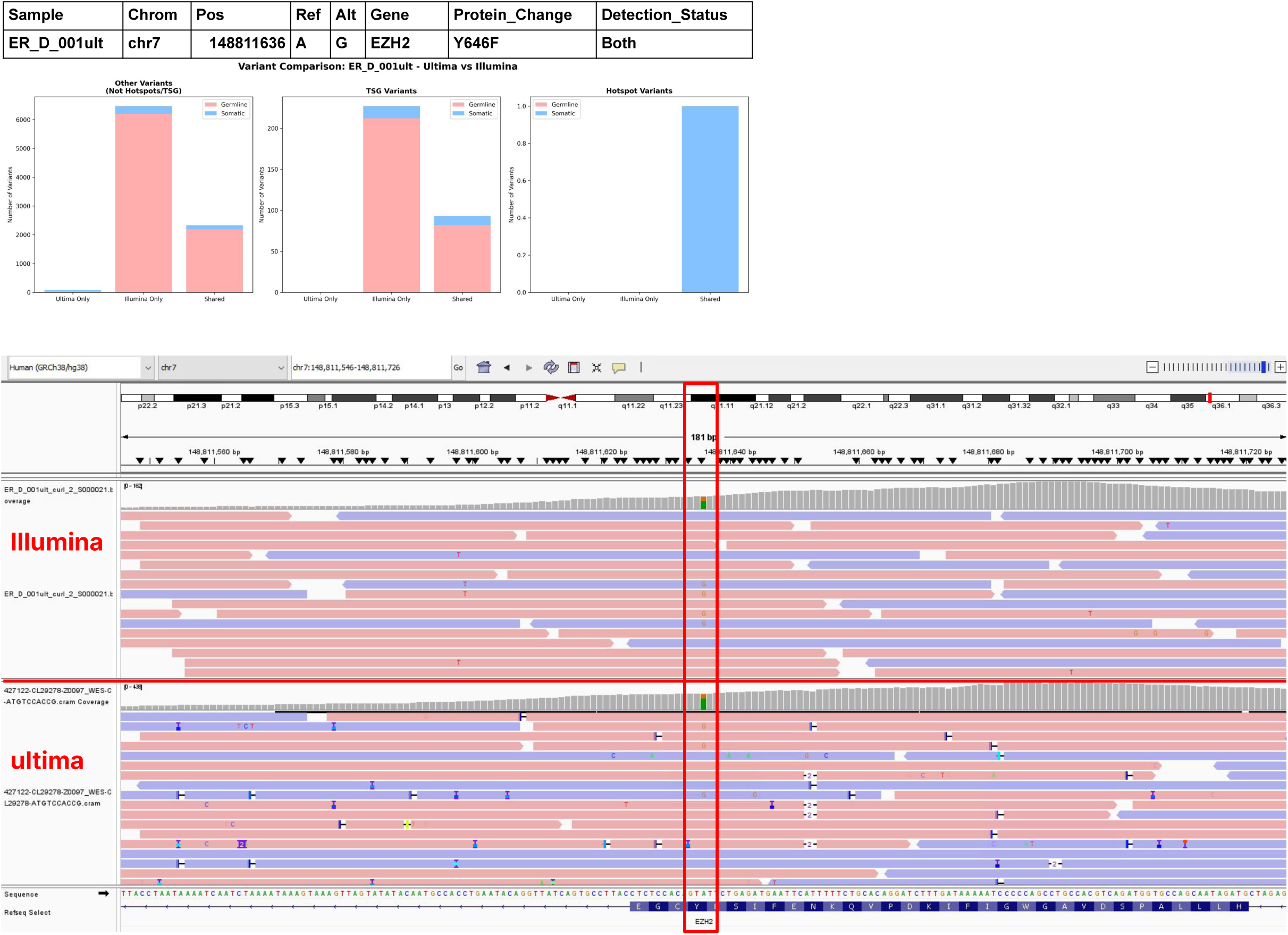
WES. Genome browser evaluation of the variant with position 148811636 in Chr7 from gene EZH2 detected in both technologies

