## Supplementary Tables for "Multi-modal benchmarking of the Ultima UG100 and Illumina NovaSeq sequencing platforms using clinically relevant FFPE tissues"

| | |
|---|---|
| Table 1 | List of the study samples and QC metrics |
| Table 2 | Unexpresed genes from each technology |
| Table 3 | List of key genes for each indication |
| Table 4 | Pathways found in both technologies |
| Table 5 | Differentially expressed transcripts between both technologies |
| Table 6 | Clinically actionable variants identification |
| Table 7 | Functional annotation WES |
| Table 8 | Variants for WGS benchmarking |
